# Longitudinal versus Cross-sectional effects of Age on MEG Power Spectra Parameters: Implications for Normative Models and Brain Ageing

**DOI:** 10.64898/2026.06.01.729181

**Authors:** Maite Crespo-Garcia, Dace Apšvalka, Ina Demetriou, Adam Attaheri, Tina Emery, Máté Aller, Richard Henson

## Abstract

Ageing produces changes in brain function. To map this process, researchers increasingly utilise Magnetoencephalography (MEG) to identify spectral shifts across large, age-diverse cohorts. However most such research has used cross-sectional designs, which may be confounded by generational differences and high inter-individual variability. To address these limitations, we analysed longitudinal resting-state MEG data from the Cam-CAN cohort, an adult lifespan cohort with a repeat MEG recording after approximately a decade. This design allowed us to contrast cross-sectional age differences between people (at baseline) with longitudinal age changes within people. Our analyses revealed four key findings. First, we observed consistency between baseline and longitudinal effects of increasing age on the slowing of the Alpha peak frequency. Second, we identified non-linear changes in Beta power, whereby power increased from youth to mid-life, but decreased from mid-life to late-life. These findings validate the use of normative charts for mapping these spectral components. Third, we detected some longitudinal effects, such as power reductions in Alpha and slowing of Beta peak frequency, that were not apparent cross-sectionally, underscoring the importance of intra-individual designs. Fourth, we did not find the commonly reported age-related flattening of the aperiodic exponent of the power spectrum, except in direct measures of cardiac activity. Together, these results demonstrate that while cross-sectional data capture major trends, longitudinal data are essential for isolating the true neurophysiological signatures of the ageing process.

## Introduction

Healthy ageing affects functional activity in the brain – as measured by MEG, for example – possibly prior to detectable effects on brain structure. The recent availability of relatively large cohorts with MEG data has led researchers to develop normative charts, which characterise, for instance, the spectral features typical of a certain age (Hoshi & Shigihara, 2020; Ranasinghe et al., 2025; Thuwal, Banerjee, & Roy, 2021; Zamanzadeh et al., 2026). While useful for characterising individual differences, such charts may be less useful for understanding the effect of ageing on the human brain. This is because they are largely informed by cross-sectional studies, in which measurements at a single time-point are compared across individuals with a different birth year. Any age-related differences between these individuals could owe to “cohort effects”, such as generational changes in diet or pollution, rather than age per se. Furthermore, other individual differences, such as head size or cortical folding, may swamp age-related differences in MEG spectral properties.

Longitudinal studies, in which changes are assessed across multiple time-points within the same individual, are therefore more informative about the ageing process. Indeed, analyses have shown that cross-sectional and longitudinal effects of age often dissociate, such as in structural brain measures (Di Biase et al., 2023). However, longitudinal studies are more difficult to run, particularly over a time-period long enough for appreciable brain changes to occur (e.g., multiple years), and there are few longitudinal studies with MEG data. Fortunately, the Cam-CAN project has recently acquired resting-state MEG data on the same individuals after a gap of up to 13.3 years (Demetriou et al., 2025). Moreover, because these adults were born across a large range of years (they were aged 18-88 at the first, baseline time-point), we can statistically compare cross-sectional age differences with longitudinal changes due to age*ing*. Similar effects (and effect sizes) for age would reinforce the utility of the normative charts based on cross-sectional data, while different effects would point to scientifically-important confounds, owing to generational changes for example. Furthermore, we can test whether the rate of within-individual changes depend on their age at baseline (i.e., non-linear effects of ageing).

Previous cross-sectional studies suggest dissociable effects of age on electrophysiological power within different frequency bands and/or within different brain regions. The most consistent finding is a slowing of the Alpha rhythm (a prominent oscillation around 8-12 Hz), which starts in late-adulthood, with the degree of slowing accelerating with age (Dustman, Shearer, & Emmerson, 1993; Klimesch, 1999). Some have distinguished low- and high-frequency power changes within the Alpha band, with widely-distributed, age-related decreases in a high-Alpha, and more posteriorly-centred, age-related increases in low-Alpha (Quinn et al., 2025). While this pattern could reflect an age-related shift of a single Alpha component towards lower frequencies, the topographical differences suggest there may be different oscillations within the Alpha band (Scally, Burke, Bunce, & Delvenne, 2018). This illustrates the importance of separating the frequency and power of such periodic components, which is why we adopted a technique to parameterise the power spectrum (Donoghue et al., 2020). This technique fits a number of Gaussians to the peaks of the power spectrum, parametrised by their peak frequency, peak amplitude and peak width.

Other notable findings include age-related differences in power in the Beta band (15-30 Hz), particularly within sensorimotor regions (Gomez, Perez-Macias, Poza, Fernandez, & Hornero, 2013; Koyama, Hirasawa, Okubo, & Karasawa, 1997; Quinn et al., 2025; Rempe et al., 2022; Rempe et al., 2023; Ruuskanen, Avendano-Diaz, Liljestrom, & Parkkonen, 2026). While some report a linear increase with age (Quinn et al., 2025; Ruuskanen et al., 2026; Shou, Yuan, Cha, Sweeney, & Ding, 2022), others claim a non-linear, “inverted-U” shape across the lifespan (Gomez et al., 2013; Rempe et al., 2023; Sahoo, Pathak, Deco, Banerjee, & Roy, 2020; Stier, Braun, & Focke, 2023), with beta power peaking around the fifth or sixth decade (Gomez et al., 2013; Ranasinghe et al., 2025; Stier et al., 2023). These trajectories might differ for Beta sub-bands, with Low-Beta showing a more linear relationship, and High-Beta showing a more quadratic relationship (Gomez et al., 2013; Stier et al., 2023). For the Gamma band (typically 30–80 Hz), some studies report a general age-related increase throughout adulthood (Quinn et al., 2025; Ruuskanen et al., 2026; Stier et al., 2023), while others suggest a decline or spectral shift in later life (Gomez et al., 2013; Rempe et al., 2023; Stier et al., 2023). At the other end of the spectrum, power in Theta (4-8 Hz) and Delta (2-4 Hz) bands have shown linear decreases in power across the lifespan (Quinn et al., 2025; Rempe et al., 2023; Shou et al., 2022; Stier et al., 2023), though again, other studies suggest a U-shaped relationship in which low-frequency power reaches a minimum during middle-age, before plateauing or increasing in later decades (Gomez et al., 2013; Rempe et al., 2023). Some of this non-linear behaviour could reflect cohort effects.

Finally, spectral parametrisation techniques also allow one to fit an “aperiodic” component, by assuming it has a “1/f” form, where power is inversely related to frequency (f). Indeed, this is methodologically important, because studies have demonstrated that the effects of age on periodic components like Alpha can be significantly over-estimated if the underlying aperiodic component is not first subtracted (Merkin et al., 2023; Trondle et al., 2023). Furthermore, effects of age on the aperiodic component itself may be scientifically important (Campbell et al., 2026; Montemurro et al., 2024). For example, the 1/f exponent has been proposed as a proxy for the excitatory-to-inhibitory (E/I) ratio in brain dynamics (Gao, Peterson, & Voytek, 2017). A common finding is an age-related ‘flattening’ of the power spectrum (i.e., smaller 1/f exponent, Voytek et al., 2015), which has been attributed to an age-related increase in the E/I ratio. Clinically, this spectral flattening has also been associated with poorer cognitive outcomes in patients with Alzheimer’s disease (van Nifterick et al., 2023). Nevertheless, other recent work has questioned such claims, by showing that many effects on 1/f slope are driven by age-related differences in cardiac activity (Schmidt et al., 2025). We therefore took care to examine and control for cardiac activity in our MEG data.

In summary, by applying spectral parametrisation to a longitudinal MEG dataset that is unique in both the length of follow-up and range of baseline ages, we could dissociate baseline (level) from change (slope) effects of age, and their potential interaction, while controlling for a range of potential confounds. The findings inform both normative (and therefore clinical) application, as well as widen scientific understanding of the effects of age on brain electrophysiology.

## Materials and methods

### Dataset and participants

Resting-state MEG data were collected on N=137 members of the Cam-CAN cohort, who completed both Phase 5 (which ran from 2024-2025) and Phase 2 (either Arm 1: 2011-2013 or Arm 2: 2016-2018; see Demetriou et al., 2025). However, four of these were excluded because: 1) one participant’s data file was corrupted; 2) one participant had a head position that was >3 SDs more anterior than the rest of the sample; 3) one participant had a head position whose Euclidean distance from the mean position was >3 SDs (in addition to being reported as restless and needing to be repositioned several times during the experiment); 4) one participant’s data showed strong magnetic interference due to metallic orthodontics. This left n=133 participants (266 recordings). Distributions of their age, sex and lag between measurements are shown in Figure 1.

**Figure 1.**
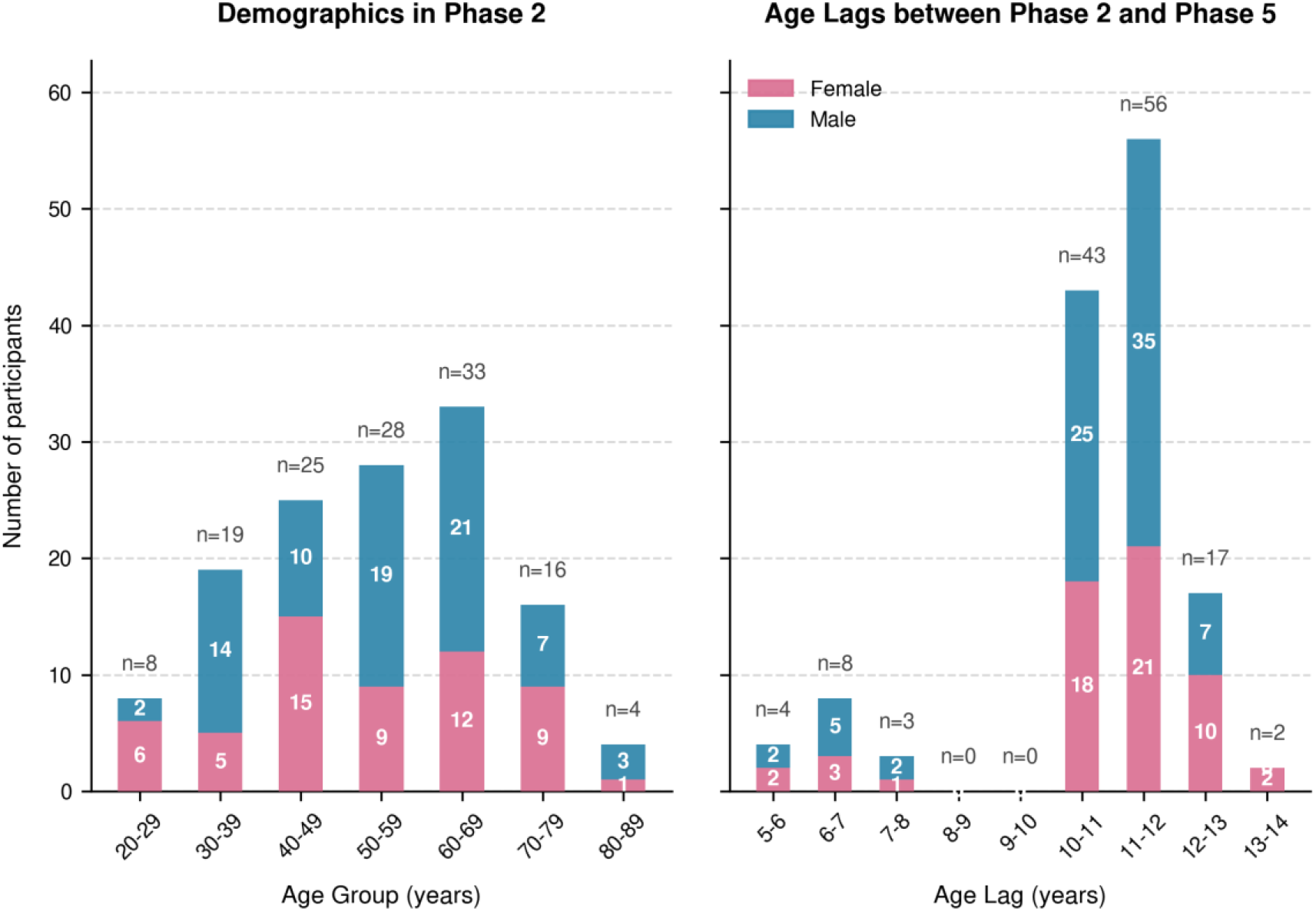
Distributions of baseline age, sex and lag in the Cam-CAN MEG longitudinal sample. (Left) Number of participants within each age group (7 decades between 20 to 90 years) at their first session (Phase 2). Stacked colour bars show the number of female (pink) versus male (blue) participants within each decade. (Right) Number of participants for each possible lag between their first and second session, rounded to nearest year, and again split by sex.

All individuals were free of self-reported neurological or psychiatric conditions (e.g. dementia, epilepsy, head injury with severe sequelae, bipolar disorder, schizophrenia), had no history of substance abuse, no communication problems (hearing, speech, or visual impairment), and no MRI/MEG contraindications (for further details, see Shafto et al., 2014). All participants scored 24 or higher on the Mini Mental State Examination (score >= 24; Folstein, Folstein, & McHugh, 1975) in both Phase 2 and Phase 5, except for three of them. Two of these were from Arm 2 of Phase 2, and one was from Arm 1 but only dropped below 24 in Phase 5. However since none had not sought clinical diagnosis, all three were included. The study was conducted in accordance with the Declaration of Helsinki (World Medical Association, 2013), and approved by the local ethics committee, Cambridgeshire Research Ethics Committee (10/H0308/50). The raw data from Phase 2 are already available on the CamCAN repository (https://cam-can.mrc-cbu.cam.ac.uk/dataset); data from Phase 5 will be added soon (and available on request until then).

### MEG acquisition

All MEG recordings were collected at the same site (MRC Cognition and Brain Science Unit, University of Cambridge, UK). However, the MEG system was upgraded between Phase 2 and Phase 5, in March 2020. Both systems had the same sensor configuration of 102 magnetometers and 204 planar gradiometers, but the Phase 2 system was a VectorView system from Elekta Neuromag, whereas the Phase 5 system was a fourth-generation TRIUX neo MEG system from MEGIN. This upgrade entailed mainly improved sensors and electronics. Because this hardware change is perfectly confounded by wave, we cannot model it statistically, so we utilised empty-room data from both waves to test for effects of scanner change across time.

Prior to entering the MEG scanner, three fiducials (nasion and left and right pre-aricular points) were digitised with a Polhemus system, the intersection of which defined the origin of head-coordinate system. The position of five head-position indicator (HPI) coils were also digitised, as well as approximately 100 head-shape points across the scalp and face.

The MEG data were sampled at 1 kHz and online filtered at 0.03–330 Hz. Participants were measured in a seated position inside a magnetically shielded room for approximately 9 min and 30 s. They were instructed to sit still and keep their eyes closed, but not fall asleep. During the recordings, the HPI coil positions within the MEG helmet were continuously monitored. Additionally, participants’ electrocardiogram (ECG) and electrooculogram (EOG, horizontal and vertical) were recorded to track cardiac signals and eye-movements.

### MEG preprocessing

MEG raw data files were organised according to the Brain Imaging Data Structure (BIDS) format (Niso et al., 2018), and preprocessed with the MNE-BIDS automatic pipeline (Jas et al., 2018) and custom scripts based on MNE-Python (Gramfort et al., 2013; Larson, 2025). These scripts are available on the GitHub repository (Crespo-García, 2026), but are summarised below (see also Supplementary Figure 1 for schematic of pipeline).

We started with the MNE-BIDS automatic pipeline. This first checked for flat and noisy channels, based on unusually low and high variability, respectively, after low-pass filtering to 40 Hz. Then, the MNE Maxwell-filter function was used to suppress environmental noise and adjust for head movement. More specifically, we applied temporal signal space separation (tSSS) with buffer duration of 10 s, correlation limit of 0.98, and internal order for the Maxwell basis of 8. The head origin parameter was set to ‘auto’ to estimate the origin of sphere fit to the headshape points, excluding those around the nose, within the coordinate system of the device. The data were corrected for head motion using the default configuration parameters, while notch filters at 50, 100 and 150 Hz were applied to reduce power-line interference. To standardise the head position across participants and sessions, we subsequently transformed the data as if the origin of the head coordinate system was in a position of (0, 0, 44) mm relative to the device origin. Finally, the pipeline down-sampled the data to 300 Hz, after high- and low-pass filtering at 0.1 Hz and 145 Hz respectively.

To detect artefacts (e.g., due to muscle activity), we used a custom script to mimic the procedure of (Stier et al., 2023), who examined cross-sectional effects of age in the full Cam-CAN Phase 2 sample. First, the Maxfiltered data prior to down-sampling were high-pass filtered at 1 Hz instead. After down-sampling to 300 Hz, the data were bandpass filtered between 110 and 140 Hz and the resulting magnitude envelope Z-scored for each channel. The Z-scored envelopes were then summed across channels, divided by the square root of the number of channels, low-pass filtered at 4 Hz to smooth out transient spikes and prevent false-positives, and finally, all samples with a Z-score above 14 were marked as bad (artefacts). Any non-bad data segments between artefacts that had a duration shorter than 200 ms were incorporated into adjacent artefacts.

We then applied independent component analysis (ICA) to detect cardiac and ocular artefacts. Here, we used a copy of the final down-sampled file from the initial automatic pipeline. First, the data were high-pass filtered at 1 Hz to facilitate ICA decomposition. Then, the data were segmented into 10-s epochs, and any epochs containing an artifact (detected in previous step) were removed, to prevent artefacts from impairing the source separation. Components showing correlations higher than 0.8 with either of the two EOG channels were classified as ocular artefacts, while those showing correlations higher than 0.4 with the ECG channel were classified as cardiac artefacts. Although other studies have typically chosen higher correlation thresholds to detect cardiac components, we decided to apply the more rigorous threshold to minimize the contribution of cardiac activity in the aperiodic (1/f) activity, which may confound age-related changes in brain activity (Schmidt et al., 2025). Finally, these components were projected out of the data, to generate a clean data file used in the spectral analyses below.

### ECG preprocessing

The ECG channel of each MEG dataset was preprocessed with a custom script. To ensure that any divergent results between the ECG and MEG data were not artefacts of preprocessing, we applied identical filtering and resampling parameters to both signals. Specifically, data were notch-filtered at 50, 100, and 150 Hz, band-pass filtered between 0.1 Hz and 145 Hz, and down-sampled to 300 Hz.

### Empty-room preprocessing

Each MEG dataset was associated to an empty room MEG recording that was acquired close in time, ranging from 8 days before to 9 days after (mean = 0.0 days, SD = 1.2). Empty room recordings were preprocessed using the same pipeline as above, except for the steps that were not applicable (e.g., *trans* option, movement correction, bad epoch detection and ICA decomposition).

### Spectral analyses

Preprocessed participant data were cropped to a common duration of 532 s, while empty-room data (which tended to be shorter) were cropped at 50 s. All data were segmented into non-overlapping 2-s epochs, and power spectra computed for each channel using Welch’s method (Welch, 1967) between 0.5 and 145 Hz, with a 2-s Hamming window, resulting in a frequency resolution of 0.5 Hz. To allow for participants settling into the resting state, the first 30 s of the MEG and ECG data (i.e., first 15 epochs) were removed. Epochs containing artefacts (as defined above) were removed before computing the average power spectrum across epochs for each channel.

To extract the periodic and aperiodic parameters of interest, we applied a widely-used approach for spectral parametrisation, implemented in an open-source Python toolbox (specparam, version 2.0), formerly known as FOOOF (“Fitting Oscillations & One-Over F”; Donoghue et al., 2020). The algorithm separates the spectral power into distinct components: one aperiodic, 1/f-like background activity (modelled as an exponential), and one or more periodic, band-limited peaks above the aperiodic background representing neural oscillations (modelled as Gaussians). The aperiodic component is often characterised by its exponent (equivalent to the sign-flipped slope of a linear fit in log–log space) and offset (y-intercept). The periodic peaks are characterised by a centre frequency, power and bandwidth. Since noise peaks can significantly affect the spectral fitting, we deliberately suppressed a focal peak of non-physiological origin at 23 Hz (that was well known to occur for some participants owing to lab set-up), using spectral interpolation between 21.9-23.9 Hz.

Spectral parametrisation was applied from 2-40 Hz to each channel. The algorithm was set to model a maximum number of 6 peaks. The peaks were limited to have a width between 1–6 Hz, a minimum height of 0.05 (in absolute units of log power, above the aperiodic fit), and a relative peak threshold of 1.5 standard deviations of the aperiodic power spectrum. These values have been commonly chosen in previous studies (e.g., Finley, Angus, van Reekum, Davidson, & Schaefer, 2022). See Supplementary Figure 2 for a distribution of peak counts per frequency bin.

The aperiodic component was modelled in *fixed* mode, assuming a single, consistent slope (exponent) within this frequency range. We also considered the possibility that the exponent was better characterised when fitting the aperiodic component with an additional *knee* parameter (Supplementary Figure 3). However, in contrast with a previous study (Zamanzadeh et al., 2026), the goodness-of-fit with the knee mode (R^2^) was worse than with the fixed mode (Supplementary Figure 4), possibly because that previous study did not model any periodic components. Indeed, the knee overlapped mainly with the Beta peak, and after allowing for a knee, the topographic distribution of the 1/f exponent seemed to resemble that of the Beta peak, rather than the more widespread effect expected (Supplementary Figure 5).

The ECG power spectrum was fitted with the same parameters, but in this case, adding a knee to the aperiodic component increased the goodness-of-fitness (Supplementary Figure 6), capturing a distinct change in the slope of the power decay around 10–20 Hz. The presence of a knee in the ECG has been reported in previous studies (e.g., Schmidt et al., 2025).

The spectral parameters of interest were the exponent of the aperiodic component, as well as the centre frequency and the log10 power of each periodic component (since the effect of age on the intercept of the aperiodic component is difficult to interpret, while effects of age on the width of periodic components were of less interest). Each peak was assigned to a frequency band if its centre frequency fell within the boundaries. These frequency bands were: Theta (4-8 Hz), Alpha (8-13 Hz), Beta (15-30 Hz) and Gamma (30-40 Hz), plus four further sub-bands: Low-Alpha (8-10 Hz), High-Alpha (10-12 Hz), Low-Beta (12-20 Hz) and High-Beta (20-30 Hz), to compare with findings by previous studies (Quinn et al., 2025; Stier et al., 2023). We then averaged each spectral parameter across channels of the same MEG sensor-type (i.e, magnetometers or planar gradiometers), after excluding any channel in which the corresponding parameter could not be obtained (e.g., no periodic peak found in a specific frequency band). This meant a total of 17 parameters per sensor-type (1 aperiodic + 2 x 8 periodic). Before applying the statistical models (see next section), we confirmed that the distribution of parameter estimates were close to Gaussian across participants and sessions.

### Statistical analyses

We employed linear, mixed-effects models (LMEs) to simultaneously investigate longitudinal and baseline age effects on each spectral parameter of interest. The basic model included three fixed effects: 1) a baseline age predictor, *A0*, defined as the participant’s age in years at Phase 2; 2) a longitudinal age predictor, *dA*, defined as the difference in years between the current session and Phase 2 (i.e., equal to 0 at Phase 2, or ranging from 5.5-13.3 years at Phase 5; see Figure 1); and 3) the interaction term (*A0*:*dA*), which can capture non-linear effects of age in which the rate of change (*dA*) depends on baseline (*A0*). Note that, apart from the change in scanner (see below), we are not assuming any “period” effects, i.e., effects of year of testing (Rohrer, 2025). A random intercept was also modelled for each participant. To control for potential confounding effects of other factors, we also fit extended models that included additional fixed effects, as described below. All fixed effects were centreed (demeaned) before modelling, while the additional covariates were scaled by their standard deviation (Z-scored).

The first extended model (the “empty-room” model) included the relevant spectral parameter from the empty-room data, to control for changes in the MEG system between Phase 2 and Phase 5 (see Supplementary Figure 8 for empty-room power spectra). For the periodic components, the spectral parametrisation step did not detect enough peaks within the bands of interest. Thus, this extended model could only be applied to power estimates, not the centre frequency estimates. For the power estimates, the empty-room power was first divided by the aperiodic component of the corresponding participant’s MEG data (before taking the log10 value). The purpose of this scaling was to obtain empty-room power values on the same scale as the power parameters obtained from spectral parametrisation of MEG data. Then, for each MEG channel, we computed the log10 transform of the empty-room scaled power within the band peaks detected in each MEG channel (peak centre frequency +/-peak width). Finally, we assigned each peak to a frequency band and averaged across all channels, to match what was described above for the MEG parameters. We also report the basic LME fit to the empty-room data in Supplementary Table 5.

The second extended model (the “cardiac” model) included the relevant spectral parameter from the ECG data for each participant and session, to control for any residual cardiac artefacts in the MEG data (despite the ICA de-noising described above). For the periodic components, the spectral parametrisation step did not detect enough peaks within the bands of interest, so we followed a similar approach as described for the empty-room model above. The main difference is that we scaled the ECG power spectrum by dividing it by the sum of the power spectrum within the fitted range (2-40 Hz). We opted to calculate the relative power spectrum for the ECG, rather than normalising by MEG’s aperiodic power, because the ECG signal uses different physical units than the MEG data, and exhibits significantly higher inter-participant scaling variability (e.g., due to differences in impedance, heart position relative to the sensors, etc.). We not only added this ECG covariate to the basic model, but also fit the basic LME to both the ECG data, and to the MEG data created from all those independent components that were associated with cardiac activity in the ICA (latter LMEs are reported in Supplementary Tables 6 and 7).

A final extended model (the “6-covariate” model) included 6 covariates: head position within the MEG helmet for 1) x-, 2) y- and 3) z-directions; 4) a measure of head movement (namely, the standard deviation of the Euclidean distance between the head position every 1 s during the recording, relative to the median head position across time points); 5) sex; and 6) the total intracranial volume (TIV) derived from running FreeSurfer v7.0 on a T1-weighted structural MRI for each participant from Phase 2 (see Demetriou et al., 2025). We also tested the effect of age on the covariates themselves, with the LMEs reported in Supplementary Table 8. Note that we did not find any significant effects of self-reported sex on any spectral parameter (all uncorrected p > .05), suggesting that males and females do not differ meaningfully in any of these MEG spectral measures.

### Other control analyses

In addition to the LME model analyses with covariates, we performed other control analyses. The first tested whether the results were affected by artefacts caused by the 21.9-23.9 Hz interpolation, by repeating the LME analysis using the basic model, but ignoring all the peaks that fell within the interpolated interval. In a second test, we refit the basic model but used the parameters extracted from power spectra without interpolation. These LMEs are reported in Supplementary Tables 9 and 10.

### Correction for multiple comparisons

In the main paper, we report statistics on the spectral parameters for the planar gradiometers, since their topographic distribution is easier to interpret than magnetometers (the statistics for magnetometers are reported in Supplementary Table 11). To control for multiple comparisons across the 17 dependent variables, we used a permutation approach to estimate the null distribution of the maximal statistic. For each of n = 10,000 permutations of participant and phase labels, a linear mixed-effects (LME) model was fit to each variable, and the maximum absolute T-statistic across all 17 tests was recorded, for each age effect (*A0, dA* and their interaction). This procedure generated an empirical null distribution of max-T values for each age effect, the 95th percentile of which defined the threshold for family-wise error (FWE) correction with α = 0.05. This approach allows for inter-correlations among the tests.

## Results

To help visualisation of the results (particularly for interactions between ageing, dA, and baseline age, A0), we divided the sample into 3 age groups based on a tertile split of baseline age at Phase 2: Young (24.3 - 47.6 years at baseline), Middle (47.8 - 63.1 years at baseline) and Old (63.2 - 84.1 years at baseline).

### Overall Power Spectra

Before reporting statistical tests on the spectral parameters, we visualised effects of cross-sectional and longitudinal age on the basic power spectra (Figure 2). The most noticeable effects were a slowing and weakening of the Alpha peak, and a reduction in power in the Beta band, for both types of age effect (note the smaller longitudinal effects are likely to reflect the smaller age range). This was more apparent after removing the aperiodic 1/f component (lower panels).

**Figure 2.**
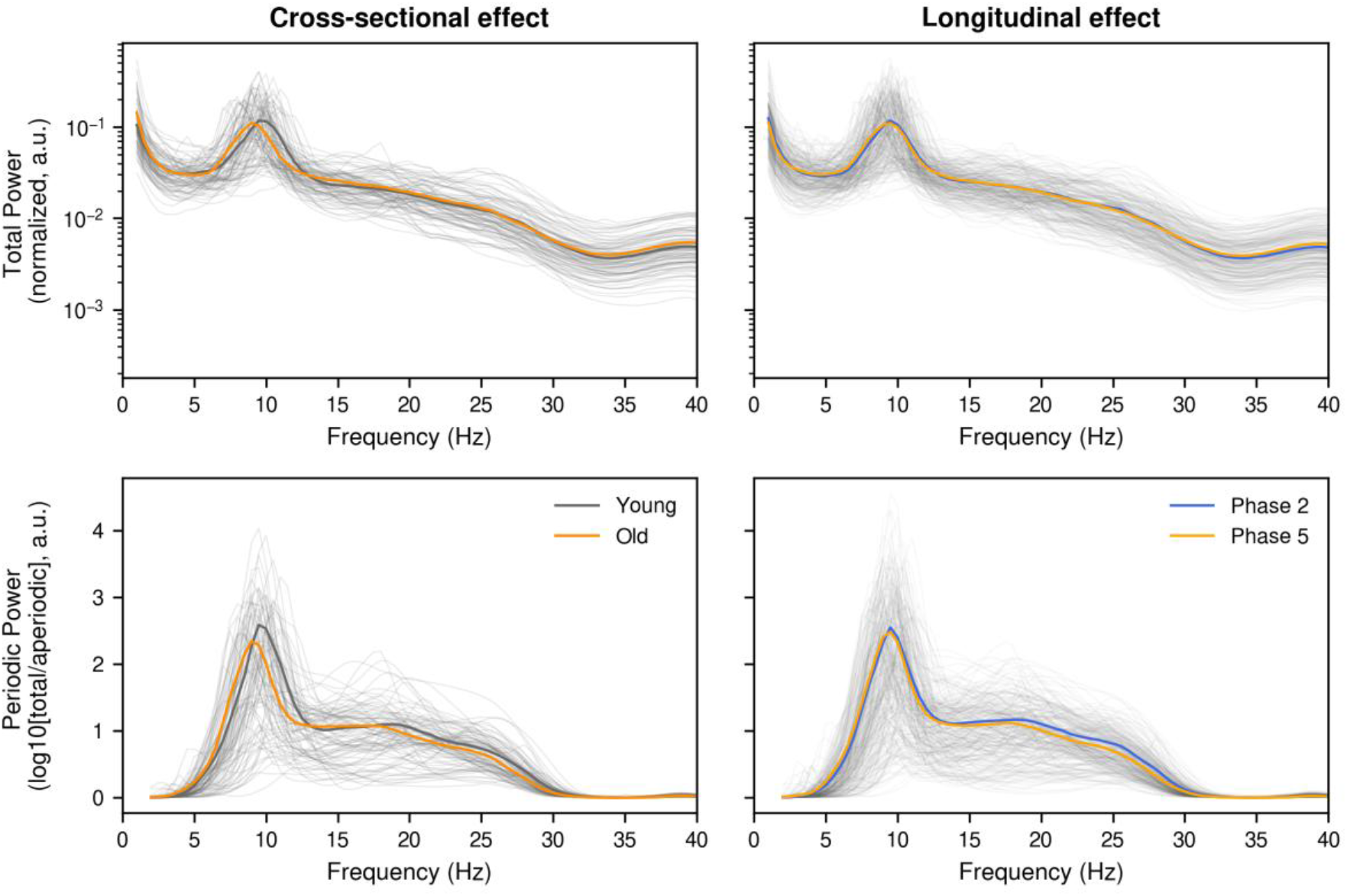
Effects of cross-sectional (A0, left) and longitudinal (dA, right) age on raw (upper) and aperiodic-adjusted (lower) power spectra. Each thin line represents an individual participant (averaged across sessions on left), while thick lines represent average across participants (separately for Young and Old groups on left, or for Phase 2 and Phase 5 on right). There is clear slowing and weakening of the Alpha peak cross-sectionally, and to a lesser extent, longitudinally (though note the cross-sectional age difference is approximately 40 years, whereas the longitudinal age differences is only approximately 12 years), which is more apparent after adjusting for the aperiodic effect. There is also a suggestion of a reduction in Beta band power (20-30 Hz).

To further examine whether the effects of longitudinal change depended on baseline age, we also plot longitudinal effects for each age group in Figure 3. These plots clearly show accelerated effects of ageing on power from 15-30 Hz (Beta band), with more pronounced decreases in the Old group relative to the other two groups. Indeed, there was a suggestion of the opposite pattern of power increases within the Young group in low-Beta (15-20 Hz).

**Figure 3.**
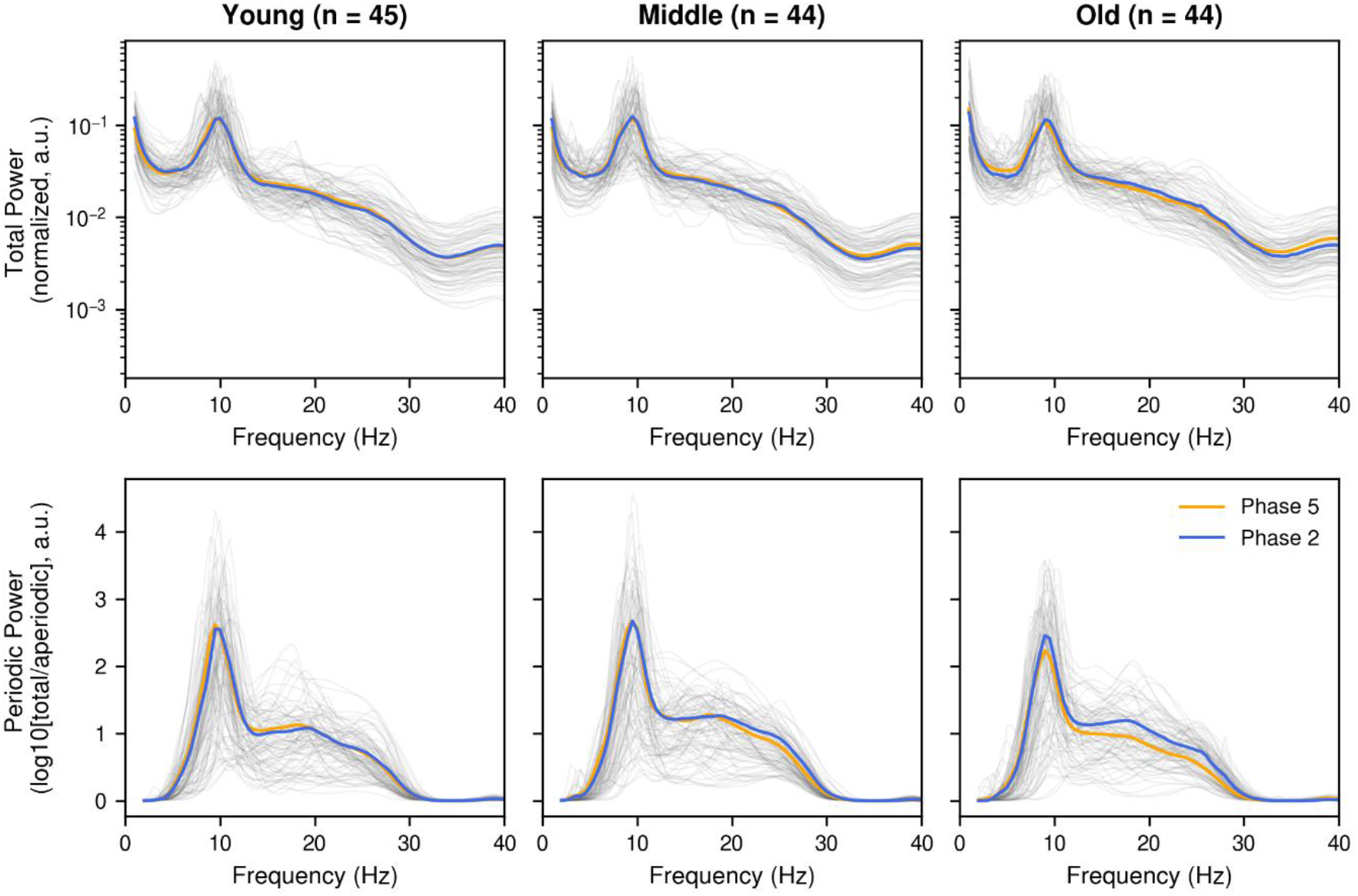
Longitudinal effects as a function of baseline age group (columns), on raw (upper) and aperiodic-adjusted (lower) power spectra. See Figure 2 legend for more details.

### Statistical Tests

Here we report statistical tests of the effects of A0 (baseline age), dA (time lag), and their interaction (A0:dA), on spectral parameters estimated from the planar gradiometers (results for the magnetometers were very similar, and shown in Supplementary Table 11). We focus on those effects that were robust across our sensitivity analyses, i.e., remaining after adjusting for 1) empty-room estimates of scanner change, 2) potential cardiac confounds, and 3) head position/motion/size and sex (see Methods), but also comment on cases where an age effect depended on such covariates (since adjusting for covariates could also remove true effects of age that happened to be correlated with those covariates).

### Aperiodic exponent

We did not find significant effects of age on the mean aperiodic 1/f exponent, even in the basic model (Table 1). This pattern did not change much when adjusting from scanner changes (empty-room model), cardiac activity (cardiac model), nor covariates related to head position/motion/size and sex (6-covariate model). In general, the exponent was largest (collapsing across all participants and sessions) around peripheral lateral and posterior sensors (top topography in middle column of Figure 4).

**Table 1.** Aperiodic (1/f) exponent statistics for the three fixed effects of baseline age (A0), change in age (dA) and their interaction (A0:dA) from the four Linear Mixed Effects (LME) models. Abbreviations: a.u. = arbitrary units, p.a. = per annum (for main effects of A0 and dA). Effects that survived correction in bold.

| Aperiodic Exponent Model / Age Effect | Parameter Estimate<br>a.u. p.a. (SD) | T-statistic (df) | P-value uncorrected (corrected) |
| --- | --- | --- | --- |
| **Basic Model** |  |  |  |
| A0 | -0.0011 (0.00091) | -1.24 (131.2) | 0.22 (0.94) |
| dA | -0.0011 (0.00098) | -1.15 (131.9) | 0.25 (0.97) |
| A0:dA | 0.0002 ($6.9 \times 10^{-5}$) | 2.91 (132.4) | 0.0043 (0.06) |
| **Empty-room Model** |  |  |  |
| A0 | -0.0063 (0.0055) | -1.15 (131.5) | 0.25 (0.97) |
| dA | -0.0056 (0.006) | -0.94 (131.7) | 0.35 (1) |
| A0:dA | 0.0012 (0.00041) | 2.88 (130.6) | 0.0047 (0.065) |
| **Cardiac Model** |  |  |  |
| A0 | -0.0057 (0.0056) | -1.02 (134.6) | 0.31 (0.99) |
| dA | -0.0061 (0.0059) | -1.04 (132.2) | 0.3 (0.99) |
| A0:dA | 0.0012 (0.00042) | 3.0 (132.1) | 0.0033 (**0.047**) |
| **6-covariate Model** |  |  |  |
| A0 | -0.0013 (0.0009) | -1.46 (129.0) | 0.15 (0.84) |
| dA | -0.00015 (0.0011) | -0.14 (135.0) | 0.89 (1) |
| A0:dA | 0.0002 ($7 \times 10^{-5}$) | 2.9 (127.0) | 0.0044 (0.063) |

**Figure 4.**
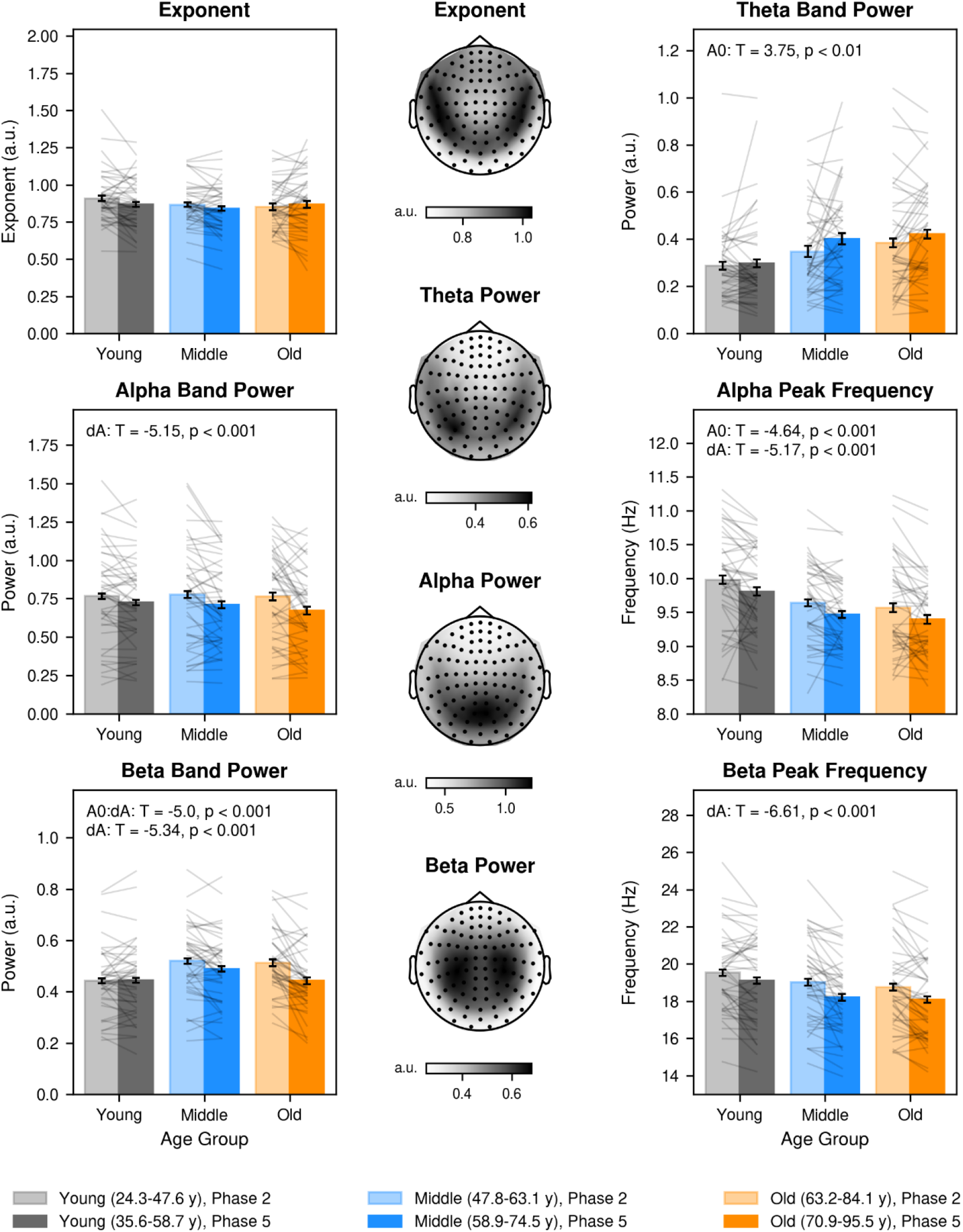
Bar plots and topographies of relevant parameters (for gradiometers). To visualize any continuous interactions, each bar represents the mean parameter value, averaged across participants within the same age group (Young, Middle or Old at baseline) for each experimental session (Phase 2 or Phase 5). Error bars represent the standard error of the difference between sessions (identical within each age group, and such that non-overlapping error bars are likely to be significant), while grey lines illustrate individual participants. Topographies represent the mean parameter across all participants and sessions (averaged across x/y gradients at each gradiometer location).

Though it did not reach significance, the effect of A0 was negative, consistent with previous claims of an age-related flattening (Donoghue et al., 2020; Voytek et al., 2015). However, it is possible that the effects in previous studies reflected age-related changes in cardiac power, since our ECG data did show a highly significant flattening with baseline age (A0: −0.031 a.u. per year, T(131) = −2.91, p-FWE < 0.01), as well as a numeric decrease with lag (dA: −0.018 a.u. per year, T(132) = −1.41, p-FWE > 0.4) (Supplementary Table 6).

Having said this, it should be noted that we did find a marginal *A0*:*dA* interaction (p-FWE = .06), which survived correction for the magnetometers (see Supplementary Table 11), and also when adjusting gradiometer data for cardiac activity (Table 1). This positive interaction reflected a smaller exponent reduction with lag (dA) as baseline age (A0) increased. This can be seen in Figure 4 (top left), where only Young and Middle groups show a decrease across sessions (flattening). This interaction effect was not significant in the ECG channel (p-FWE > 0.4), nor in the MEG data derived from cardiac independent components (p-FWE = 1; see Supplementary Tables 6 and 7).

Given that previous studies have reported an effect of age on 1/f exponent, we re-analyzed the MEG data without removing cardiac components, in case these components had removed true neural activity too. The effects of age on the aperiodic exponent remained non-significant (A0: uncorrected p = 0.22, dA: uncorrected p = 0.15, A0:dA: uncorrected p = 0.0047, but latter did not survive correction). This suggests that while cardiac contamination could potentially bias the aperiodic exponent toward the age-related flattening reported in the literature, its removal does not seem to account for the lack of age effects in our sample.

### Theta band

Contrasting with previous studies, Theta band power significantly increased with baseline age (A0), as shown in Table 2. Its topographic distribution (middle panel of Figure 4) was similar to the aperiodic exponent, though somewhat more clustered around left and right temporoparietal sensors. It also increased from baseline to follow-up session (dA), though only in gradiometers, and not when controlling for other covariates. Any interaction between A0 and dA did not reach significance.

**Table 2.** Theta power statistics for the three fixed effects of baseline age (A0), change in age (dA) and their interaction (A0:dA) from the four Linear Mixed Effects (LME) models. Abbreviations: a.u. = arbitrary units, p.a. = per annum (for main effects of A0 and dA). Effects that survived correction in bold.

| Theta Power Model / Age Effect | Parameter Estimate<br>a.u. p.a. (SD) | T-statistic (df) | P-value uncorrected (corrected) |
| --- | --- | --- | --- |
| **Basic Model** |  |  |  |
| A0 | 0.004 (0.0011) | 3.75 (131.2) | 0.00027 (**0.003**) |
| dA | 0.0033 (0.0011) | 3.1 (131.8) | 0.0024 (**0.038**) |
| A0:dA | $6.3 \times 10^{-5}$ ($7.5 \times 10^{-5}$) | 0.84 (132.2) | 0.4 (1) |
| **Empty-room Model** |  |  |  |
| A0 | 0.019 (0.0053) | 3.67 (130.5) | 0.00035 (**0.0042**) |
| dA | 0.014 (0.0055) | 2.62 (137.1) | 0.0099 (0.14) |
| A0:dA | 0.00027 (0.00038) | 0.72 (131.0) | 0.47 (1) |
| **Cardiac Model** |  |  |  |
| A0 | 0.02 (0.0054) | 3.72 (130.3) | 0.0003 (**0.0033**) |
| dA | 0.015 (0.0053) | 2.89 (132.0) | 0.0045 (0.068) |
| A0:dA | 0.00035 (0.00037) | 0.94 (131.4) | 0.35 (1) |
| **6-covariate Model** |  |  |  |
| A0 | 0.0039 (0.0011) | 3.61 (130.6) | 0.00043 (**0.0051**) |
| dA | 0.0031 (0.0012) | 2.6 (135.1) | 0.01 (0.15) |
| A0:dA | $4.9 \times 10^{-5}$ ($7.5 \times 10^{-5}$) | 0.66 (128.5) | 0.51 (1) |

Having said this, closer inspection of the power spectra suggested that the increase in Theta power might be caused by slowing of the Alpha peak (see Figure 2-3, and next section). It is possible that the spectral parametrisation method had difficulty detecting Theta peaks, prioritizing larger, high-frequency peaks arising from the (side of) the Alpha component. This is consistent with nearly a third of datasets producing a peak estimate at the maximum of the Theta band (7 Hz) (see also Supplementary Figure 2 for a distribution of peak counts per frequency bin). Thus, these effects on Theta power should be interpreted with caution (as should those in other studies that restrict analyses to such bands).

We did not find any significant effects on Theta peak frequency.

### Alpha band

In terms of power, Alpha decreased with ageing, with a highly significant effect of dA, regardless of any adjustments (left columns of Table 3), while any effect of A0 or interaction failed to reach significance (Figure 4, middle left). It had a more central, posterior distribution than Theta (Figure 4, middle panel).

**Table 3.** Alpha statistics for the three fixed effects of baseline age (A0), change in age (dA) and their interaction (A0:dA) from the four Linear Mixed Effects (LME) models. Abbreviations: a.u. = arbitrary units, p.a. = per annum (for main effects of A0 and dA). Effects that survived correction in bold.

| Model / Age Effect | Alpha Power |  |  | Alpha Frequency |  |  |
| --- | --- | --- | --- | --- | --- | --- |
|  | Parameter Estimate, a.u. p.a. (SD) | T-statistic (df) | P-value uncorrected (corrected) | Parameter Estimate, Hz p.a. (SD) | T-statistic (df) | P-value uncorrected (corrected) |
| **Basic Model** |  |  |  |  |  |  |
| A0 | -0.0013 (0.0017) | -0.79 (131.1) | 0.43 (1) | -0.016 (0.0034) | -4.64 (131.2) | $8.2 \times 10^{-6}$ (**0.0001**) |
| dA | -0.006 (0.0012) | -5.15 (131.4) | $9.4 \times 10^{-7}$ (**< 10<sup>-4</sup>**) | -0.016 (0.0031) | -5.17 (131.7) | $8.5 \times 10^{-7}$ (**< 10<sup>-4</sup>**) |
| A0:dA | -0.00018 ($8.3 \times 10^{-5}$) | -2.12 (131.6) | 0.036 (0.39) | $-3.3 \times 10^{-5}$ (0.00022) | -0.15 (132.0) | 0.88 (1) |
| **Empty-room Model** |  |  |  |  |  |  |
| A0 | -0.0047 (0.0057) | -0.82 (128.1) | 0.41 (1) | -0.016 (0.0034) | -4.64 (131.2) | $8.2 \times 10^{-6}$ (**0.0001**) |
| dA | -0.021 (0.0041) | -5.12 (127.7) | $1.1 \times 10^{-6}$ (**< 10<sup>-4</sup>**) | -0.016 (0.0031) | -5.17 (131.7) | $8.5 \times 10^{-7}$ (**< 10<sup>-4</sup>**) |
| A0:dA | $-6.2 \times 10^{-4}$ ($2.9 \times 10^{-4}$) | -2.15 (128.0) | 0.034 (0.38) | $-3.3 \times 10^{-5}$ ($2.2 \times 10^{-4}$) | -0.15 (132.0) | 0.88 (1) |
| **Cardiac Model** |  |  |  |  |  |  |
| A0 | -0.0036 (0.0057) | -0.63 (132.9) | 0.53 (1) | -0.016 (0.0034) | -4.64 (131.2) | $8.2 \times 10^{-6}$ (**0.0001**) |
| dA | -0.021 (0.004) | -5.12 (130.7) | $1.1 \times 10^{-6}$ (**< 10<sup>-4</sup>**) | -0.016 (0.0031) | -5.17 (131.7) | $8.5 \times 10^{-7}$ (**< 10<sup>-4</sup>**) |
| A0:dA | -0.0006 (0.00028) | -2.11 (130.9) | 0.037 (0.4) | $-3.3 \times 10^{-5}$ (0.00022) | -0.15 (132.0) | 0.88 (1) |
| **6-covariate Model** |  |  |  |  |  |  |
| A0 | -0.0011 (0.0016) | -0.67 (130.0) | 0.5 (1) | -0.015 (0.0033) | -4.42 (129.5) | $2.1 \times 10^{-5}$ (**0.0003**) |
| dA | -0.0069 (0.0013) | -5.34 (131.6) | $4 \times 10^{-7}$ (**< 10<sup>-4</sup>**) | -0.02 (0.0036) | -5.47 (133.7) | $2.1 \times 10^{-7}$ (**< 10<sup>-4</sup>**) |
| A0:dA | -0.00017 ($8.1 \times 10^{-5}$) | -2.15 (127.7) | 0.034 (0.38) | $-4.3 \times 10^{-5}$ (0.00022) | -0.19 (127.3) | 0.85 (1) |

Note that, unlike previous comparisons of power at each frequency or frequency band in the power spectrum (e.g., Quinn et al., 2025; Stier et al., 2023), the present measure of power was adjusted for any concomitant change in the frequency of the Alpha peak. Indeed, that peak frequency also showed significant decreases with dA (right columns of Table 3). This time, the same was true for A0, and the interaction did not reach significance, reflecting a pattern of similar-size ageing-related reductions (slowing) regardless of baseline age (Figure 4, middle right).

Given prior claims for potentially dissociable alpha rhythms (Quinn et al., 2025), we repeated the above analyses for low and high Alpha sub-bands (Supplementary Tables 1 and 2). High-Alpha power showed the same decreases with dA, though no effects reached significance for Low-Alpha power. Nevertheless, Low-Alpha peak frequency derived from magnetometers did show a significant slowing with A0 (p-FWE = 0.021). Nonetheless, the power distributions were topographically similar (Supplementary Figure 7, upper row), providing only limited overall support for two age-related alpha effects (Quinn et al., 2025).

### Beta band

Beta power showed a significant and robust interaction between A0 and dA (Table 4), whereby power increased towards middle age, but decreased in late age (Figure 4, bottom left). This power had a bilateral central distribution. The Beta peak frequency decreased more linearly with age (Figure 4, bottom right), with a significant effect of dA, a trend in A0, and no evidence for an interaction. This pattern remained even when allowing for a sporadic artefact within the range 21.9-23.9 Hz (see Methods and Supplementary Tables 9-10).

**Table 4.**
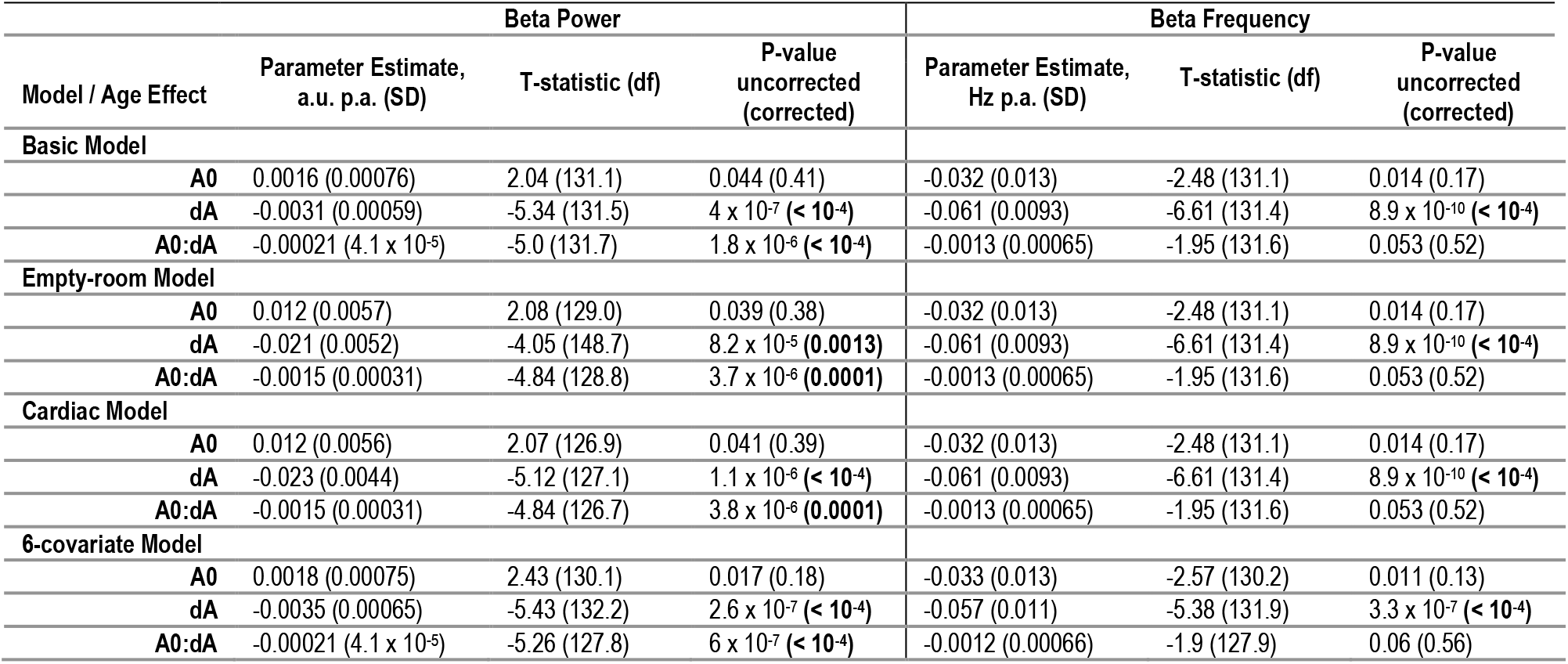
Beta statistics for the three fixed effects of baseline age (A0), change in age (dA) and their interaction (A0:dA) from the four Linear Mixed Effects (LME) models. Abbreviations: a.u. = arbitrary units, p.a. = per annum (for main effects of A0 and dA). Effects that survived correction in bold.

The interaction effect on power was evident in both High and Low sub-bands (Supplementary Tables 3-4). However, the High-Beta effect was twice the magnitude of the Low-Beta effect, and the mean band power showed a more frontal distribution than the slower sub-band (Supplementary Figure 7, bottom row). Additionally, there was a slowing of the peak frequency in both Beta sub-bands that survived correction in the High-Beta case for all models except the 6-covariate model.

### Gamma band

The spectral parametrisation algorithm only found a few Gamma peaks (in 4-62 channels, mean = 27.3), possibly because the upper limit of the fitted frequency range was 40 Hz and probably explaining why few age effects survived correction. Indeed, the only effect was an increase in Gamma peak frequency with dA, though this no longer survived correction in the 6-covariate model (Table 5) and was not present in magnetometers (Supplementary Table 11).

**Table 5.** Gamma statistics for the three fixed effects of baseline age (A0), change in age (dA) and their interaction (A0:dA) from the four Linear Mixed Effects (LME) models. Abbreviations: a.u. = arbitrary units, p.a. = per annum (for main effects of A0 and dA). Effects that survived correction in bold.

| Model / Age Effect | Gamma Power |  |  | Gamma Frequency |  |  |
| --- | --- | --- | --- | --- | --- | --- |
|  | Parameter Estimate, a.u. p.a. (SD) | T-statistic (df) | P-value uncorrected (corrected) | Parameter Estimate, Hz p.a. (SD) | T-statistic (df) | P-value uncorrected (corrected) |
| **Basic Model** |  |  |  |  |  |  |
| A0 | $7.6 \times 10^{-5}$ (0.00013) | 0.61 (130.9) | 0.54 (1) | -0.0036 (0.0063) | -0.56 (131.7) | 0.57 (1) |
| dA | -0.00058 (0.00025) | -2.34 (133.9) | 0.021 (0.26) | 0.047 (0.015) | 3.22 (136.0) | 0.0016 (**0.025**) |
| A0:dA | $1.1 \times 10^{-5}$ ($1.7 \times 10^{-5}$) | 0.65 (134.6) | 0.52 (1) | 0.0014 (0.001) | 1.34 (137.2) | 0.18 (0.92) |
| **Empty-room Model** |  |  |  |  |  |  |
| A0 | 0.00057 (0.0049) | 0.12 (141.5) | 0.91 (1) | -0.0036 (0.0063) | -0.56 (131.7) | 0.57 (1) |
| dA | -0.026 (0.0096) | -2.72 (140.7) | 0.0073 (0.11) | 0.047 (0.015) | 3.22 (136.0) | 0.0016 (**0.026**) |
| A0:dA | 0.00031 (0.00066) | 0.47 (136.0) | 0.64 (1) | 0.0014 (0.001) | 1.34 (137.2) | 0.18 (0.92) |
| **Cardiac Model** |  |  |  |  |  |  |
| A0 | 0.0027 (0.0048) | 0.57 (130.6) | 0.57 (1) | -0.0036 (0.0063) | -0.56 (131.7) | 0.57 (1) |
| dA | -0.021 (0.0096) | -2.14 (141.8) | 0.034 (0.39) | 0.047 (0.015) | 3.22 (136.0) | 0.0016 (**0.026**) |
| A0:dA | 0.00047 (0.00066) | 0.7 (135.5) | 0.48 (1) | 0.0014 (0.001) | 1.34 (137.2) | 0.18 (0.92) |
| **6-covariate Model** |  |  |  |  |  |  |
| A0 | $3.8 \times 10^{-5}$ (0.00013) | 0.29 (132.1) | 0.77 (1) | -0.003 (0.0066) | -0.45 (126.9) | 0.65 (1) |
| dA | -0.00047 (0.00027) | -1.75 (147.6) | 0.082 (0.67) | 0.041 (0.016) | 2.66 (145.0) | 0.0087 (0.12) |
| A0:dA | $1.2 \times 10^{-5}$ ($1.7 \times 10^{-5}$) | 0.71 (132.7) | 0.48 (1) | 0.0014 (0.001) | 1.36 (129.1) | 0.18 (0.91) |

## Discussion

Our analyses of the unique Cam-CAN longitudinal MEG sample allowed us to contrast cross-sectional and longitudinal effects of age on resting-state spectral parameters. First, we found consistent effects of longitudinal and baseline age on the slowing of Alpha peak frequency. Second, we found non-linear effects in Beta power changes, expressed here as an interaction between within-individual changes and baseline age. These findings support the use of normative charts to map age-normed changes in brain functional activity (to which individuals, e.g., patients, can be compared). However, we also identified unique longitudinal effects, such as power reductions in Alpha and slowing of Beta peak frequency, underscoring the importance of intra-individual designs for studying age-related changes. Additionally, we found evidence for a flattening of aperiodic exponent (reported in prior cross-sectional analyses) in direct cardiac measurements but not consistently in brain activity.

We observed the well-stablished slowing of Alpha peak frequency with advancing age (Dustman et al., 1993; Klimesch, 1999; Quinn et al., 2025; Scally et al., 2018; Stier et al., 2023), now within as well as across individuals, and robust to controlling for multiple potential confounds. Notably, the effect sizes for cross-sectional and longitudinal effects were identical, and stable across sensitivity analyses, at −0.16 Hz per decade (with a medium cross-sectional effect size of Cohen’s f^2^ = 0.16, and longitudinal Cohen’s f^2^ = 0.20). This consistency underscores the robustness of Alpha slowing as a biomarker of human ageing. We also found a decrease in Alpha power of −0.06dB to −0.21dB (relative to 1/f power) per decade within individuals (depending on covariates; Cohen’s f^2^ = 0.20-0.21). In this case however, the decrease did not reach significance for the cross-sectional effect of age (and was 1/3 to a 1/6 smaller). While this lack of cross-sectional effect is consistent with some previous studies (e.g., Ruuskanen et al., 2026; Sahoo et al., 2020), others reported age-related decreases in Alpha power (Babiloni et al., 2006; Gomez et al., 2013; Scally et al., 2018; Thuwal et al., 2021; Trubshaw et al., 2026), and yet others reported power increases (Ranasinghe et al., 2025; Rempe et al., 2023; Shou et al., 2022; Stier et al., 2023), or a coexistence of both decreases and increases in different Alpha sub-bands (Quinn et al., 2025); in some cases, even when using the same Cam-CAN cohort as used here (Quinn et al., 2025; Stier et al., 2023).

One reason for these discrepancies may be that many studies did not adjust for aperiodic power, as done here (Donoghue et al., 2020; Trondle et al., 2023). While any effect of age on the 1/f exponent did not reach significance in the present study, numerical differences in this exponent, and/or in the 1/f intercept, may have contributed to effects of age on Alpha power in previous studies. Another reason could be that many studies only examined power changes at each frequency (or frequency band), which cannot distinguish between changes in peak power and changes in peak frequency (Scally et al., 2018). Thus previous reports of age-related power decreases in high-Alpha power but increases in low-alpha (Quinn et al., 2025), could simply reflect a shift in the Alpha peak towards lower frequencies. When we separated peaks found in low (8-10 Hz) versus high (10-12 Hz) Alpha sub-bands, we found a within-individual, age-related power decrease in the high Alpha but not low Alpha sub-band. This is despite allowing for simultaneous age-related changes in peak frequency, so could indicate separate Alpha effects, though the topographic distributions provided little support for distinct high and low Alpha sources.

More generally, differences between studies are complicated by use of relative versus absolute power: while absolute MEG power at the sensors will be influenced by other factors such as head position and possibly head size (Quinn et al., 2025), relative power changes (e.g., normalised by total power across a frequency range) can miss true age-related changes in absolute power. While source space analyses (which incorporate a model of head position and size) can address such confounding factors to some extent, they necessarily require making additional assumptions about the nature of source activity (given the indeterminacy of this inverse problem). Here, we chose instead to interpolate sensor data as if (the centre of) everyone’s head were in the same position, and adjust for head-size (as well as head position and overall head motion) in our sensitivity analyses. Nonetheless, ageing also affects the thickness and curvature of the cortical surface (Magnotta et al., 1999), which will affect the MEG signal beyond head position and size. Future work could use structural MRIs that allow for such age-related cortical changes (even within-individuals) and test whether age-related MEG effects are a correlate of the same grey-matter changes, or reflect additional/independent changes in brain function. Regardless, these issues are unlikely to affect the peak frequencies of components – particularly in within-participant designs – which reinforces their value as a robust measure of ageing.

Our study also revealed an effect of age that is reported less frequently: a slowing of the Beta peak frequency (Brady & Bardouille, 2022; Karvat, Crespo-García, Vishne, Anderson, & Landau, 2026). Notably, this effect was only evident within individuals, further reinforcing the importance of longitudinal designs to uncover subtle neurophysiological changes associated with the ageing process. A linear beta slowing with age, across participants, has been observed during motor tasks in other populations (Rossiter, Davis, Clark, Boudrias, & Ward, 2014). Notwithstanding the above difficulties of interpreting power changes, we also identified a non-linear relationship between Beta power and age, where power increased with age in young individuals, but decreased with age in older individuals. This resembles the inverted-U shape described in some cross-sectional studies, where Beta power peaks between the fifth and sixth decades of life (Gomez et al., 2013; Ranasinghe et al., 2025; Stier et al., 2023). By demonstrating that these late-life declines occur within the same individuals over time, our study confirms that these patterns represent true ageing processes, rather than mere generational differences.

An unexpected finding was the lack of age-related effects on the aperiodic exponent. This stands in stark contrast to the literature, which consistently reports a flattening of this component with age (Finley et al., 2024; Finley et al., 2022; Leroy, Bublitz, von Dincklage, Antonenko, & Fleischmann, 2025; McKeown et al., 2025; Merkin et al., 2023; Montemurro et al., 2024; Ranasinghe et al., 2025; Trondle et al., 2023; Trubshaw et al., 2026; Voytek et al., 2015). While some of this divergence may stem from methodological variations, one likely contribution may derive from our artifact rejection strategy. Specifically, we employed a liberal correlation threshold (r > 0.40) to identify and remove components correlated with cardiac activity (as measured by ECG), following recent claims that age-related differences in the 1/f slope are driven by cardiac interference (Schmidt et al., 2025). Indeed, we found strong effects of age on the 1/f exponent of the ECG. Thus, it is possible that age does not affect the 1/f exponent of brain activity, and previous reports of this reflect contamination of by age-related cardiac changes.

### Caveats

Our study is subject to several methodological considerations. First, while we argue above that spectral parametrisation offers significant advantages over traditional band-wise analysis, it can be sensitive to specific parameters (such as number of peaks) and assumptions (e.g., Gaussian-shaped peaks). It may also conflate overlapping components, or fail to detect peaks in the presence of high noise levels. The latter is relevant to our Theta and Gamma bands, which exhibited a lower detection count than Alpha and Beta. The former is pertinent to Theta power, where our failure to detect the age-related reductions reported in the literature (Quinn et al., 2025; Rempe et al., 2023; Shou et al., 2022; Stier et al., 2023) may reflect mis-identification of the side-lobes of a strong Alpha peak as Theta peaks (see Results). Some of these methodological issues may be addressed by recent Bayesian extensions of spectral parametrisation (Medrano, Alexander, Seymour, & Zeidman, 2025).

Secondly, our analysis was conducted on averaged sensor-level spectral parameters. While this averaging may increase signal-to-noise ratio, it cannot resolve spatially distinct patterns that overlap in frequency. Thirdly, our longitudinal effects were confounded by a change in MEG scanner, and while we attempted to correct for this (at each frequency) using empty-room data, it is possible that this correction was insufficient, at least for the main effect of dA.

Finally, some of the present findings may be specific to the Cam-CAN sample. While participants were likely to be more representative than those in many previous studies (given our opt-out, GP-based recruitment, rather than more typical adverts; see Shafto et al., 2014) for more details), and were approximately matched for sex, they were not racially nor culturally diverse. Moreover, there are likely to be selection biases, particularly in the older group, such as those healthy enough to undergo MEG and MRI scanning, and those who returned for Phase 5 a decade after Phase 2. Thus, future studies should seek to replicate our results in a wider range of more diverse individuals.

### Other Future Work

In addition to using other methodological techniques and replicating in other cohorts, one future extension would be to distinguish rhythmic from arrhythmic activity. While peaks like Alpha and Beta are often assumed to be oscillatory, they could arise from transient, non-oscillatory impulses (bursting). Recently-developed algorithms can distinguish rhythmic from arrhythmic activity, which can in turn be used to define frequency bands from the data, rather than using a priori ranges (Karvat et al., 2026). Other analyses could take a multivariate approach: rather than trying to test effects of age on independent spectral features (the univariate approach used here), one could use multivariate techniques like tensor component analysis (Calmus, Demetriou, Crespo-García, & Henson, 2026), or partial least squares, in order to identify age-related patterns across frequency (and possibly across space). One could also examine whether age-effects generalise beyond the resting-state, by using task-based or movie-related data. Finally, one could relate these spectral parameters to other age-related changes, such as in brain structure (as discussed earlier) and/or cognitive performance.

## Data and Code Availability

The raw data from Phase 2 are already available on the CamCAN repository (https://cam-can.mrc-cbu.cam.ac.uk/dataset); data from Phase 5 will be added soon (and available on request until then).

Code is available on GitHub: https://github.com/MaiteCG/CamCAN_MEGLongRest_SpecparamLME_2026/

## Author Contributions

M.C.G.: methodology (data processing pipeline), software (preprocessing and analysis pipelines), formal analysis, writing (original draft, review and editing), and visualisation. D.A.: software (conversion into BIDS format), data curation, writing (review and editing). I.D.: investigation, writing (review and editing). A.A.: data curation, writing (review and editing). T.E.: investigation, data curation, writing (review and editing). M.A.: software (scripts and support for using MNE-BIDS and MNE-BIDS-pipeline), writing (review and editing). R.H.: conceptualisation, writing (original draft, review and editing), supervision, project administration and funding acquisition.

## Ethics

The study was conducted in accordance with the Declaration of Helsinki (World Medical Association, 2013), and approved by the local ethics committee, Cambridgeshire Research Ethics Committee (10/H0308/50).

## Acknowledgments

We thank all the Cam-CAN participants for their time and interest. We also thank the Cam-CAN corporate author, which also includes: Lorraine K Tyler, Carol Brayne, Edward T Bullmore, Andrew C Calder, Rhodri Cusack, Tim Dalgleish, John Duncan, Fiona E Matthews, William D Marslen-Wilson, James B Rowe, Meredith A Shafto; Karen Campbell, Teresa Cheung, Simon Davis, Linda Geerligs, Rogier Kievit, Anna McCarrey, Abdur Mustafa, Darren Price, David Samu, Jason R Taylor, Matthias Treder, Janna van Belle, Nitin Williams, Daniel Mitchell, Simon Fisher, Else Eising, Ethan Knights, Lauren Bates, Sharon Erzinçlioğlu, Andrew Gadie, Sofia Gerbase, Stanimira Georgieva, Claire Hanley, Beth Parkin, David Troy, Tibor Auer, Lu Gao, Emma Green, Rafael Henriques; Jodie Allen, Gillian Amery, Liana Amunts, Anne Barcroft, Amanda Castle, Cheryl Dias, Jonathan Dowrick, Melissa Fair, Hayley Fisher, Anna Goulding, Adarsh Grewal, Geoff Hale, Andrew Hilton, Frances Johnson, Patricia Johnston, Thea Kavanagh-Williamson, Magdalena Kwasniewska, Alison McMinn, Kim Norman, Jessica Penrose, Fiona Roby, Diane Rowland, John Sargeant, Maggie Squire, Beth Stevens, Aldabra Stoddart, Cheryl Stone, Tracy Thompson, Ozlem Yazlik; Dan Barnes, Marie Dixon, Jaya Hillman, Joanne Mitchell, Laura Villis.

## Supplementary Materials

Supplementary material for the current version of this manuscript is at the end of this document.

