## Supplemental Material for "Longitudinal versus Cross-sectional effects of Age on MEG Power Spectra Parameters: Implications for Normative Models and Brain Ageing"

### Supplementary Methods

#### Mean head positions of participants

The mean head position (SD) across all 266 datasets from the final longitudinal sample (N=133, after four exclusions; 2 sessions), in the MEG device space, was x = +0.28 mm (2.86), y = +7.62 mm (6.17), z = -2.79 mm (6.87).

#### MEG preprocessing pipeline

Supplementary Figure 1. Flow chart for the preprocessing pipeline applied to the Cam-CAN MEG longitudinal dataset. The dataset was initially preprocessed with the automatic MNE-BIDS pipeline (top left box) to apply Maxwell filter and frequency filters. Then, we applied other custom preprocessing steps to detect bad epochs, perform ICA and remove EOG- and ECG-related activity. Finally, we computed the power spectra that was parametrised to extract the spectral features of interest. See MEG preprocessing under Methods section in the main text.

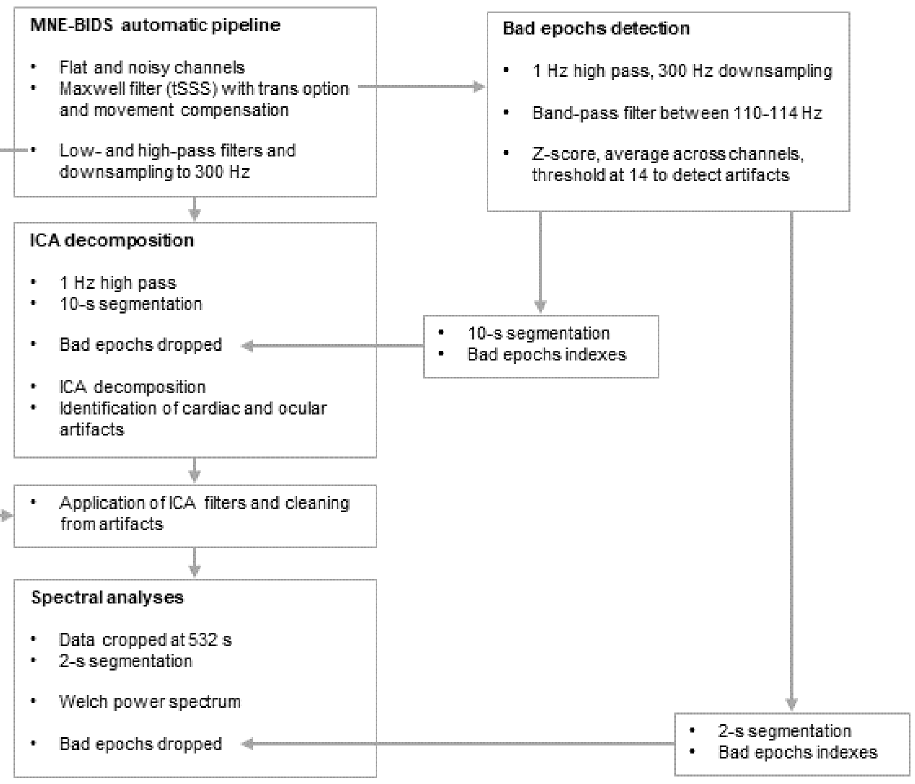


**Supplementary Figure 1. Flow chart for the preprocessing pipeline applied to the Cam-CAN MEG longitudinal dataset.** The dataset was initially preprocessed with the automatic MNE-BIDS pipeline (top left box) to apply Maxwell filter and frequency filters. Then, we applied other custom preprocessing steps to detect bad epochs, perform ICA and remove EOG- and ECG-related activity. Finally, we computed the power spectra that was parametrised to extract the spectral features of interest. See MEG preprocessing under Methods section in the main text.

#### Spectral parametrisation

Supplementary Figure 2. Number of spectral parameters detected across the whole dataset. The plot shows that the spectral parametrisation algorithm detected fewer periodic peaks within Theta and Gamma bands than in Alpha and Beta bands. Most of the Theta peaks had centre frequencies in higher frequency bins, near the boundary with Alpha.

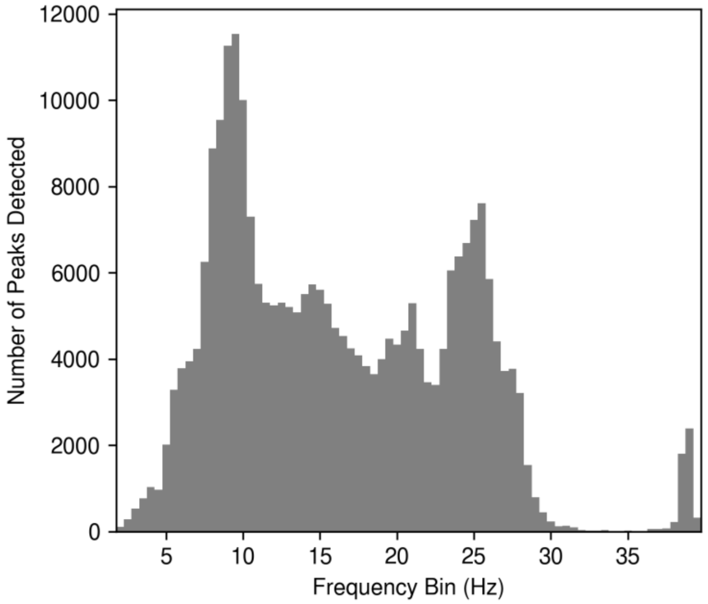


**Supplementary Figure 2. Number of spectral parameters detected across the whole dataset.** The plot shows that the spectral parametrisation algorithm detected fewer periodic peaks within Theta and Gamma bands than in Alpha and Beta bands. Most of the Theta peaks had centre frequencies in higher frequency bins, near the boundary with Alpha.

Supplementary Figure 3. Comparison of MEG aperiodic power spectrum with and without the knee parameter. The curve of the aperiodic spectrum with knee (teal) is closer to the total spectrum (black) than the fixed-mode aperiodic spectrum (brown), but this may be through also capturing the beta band peak.

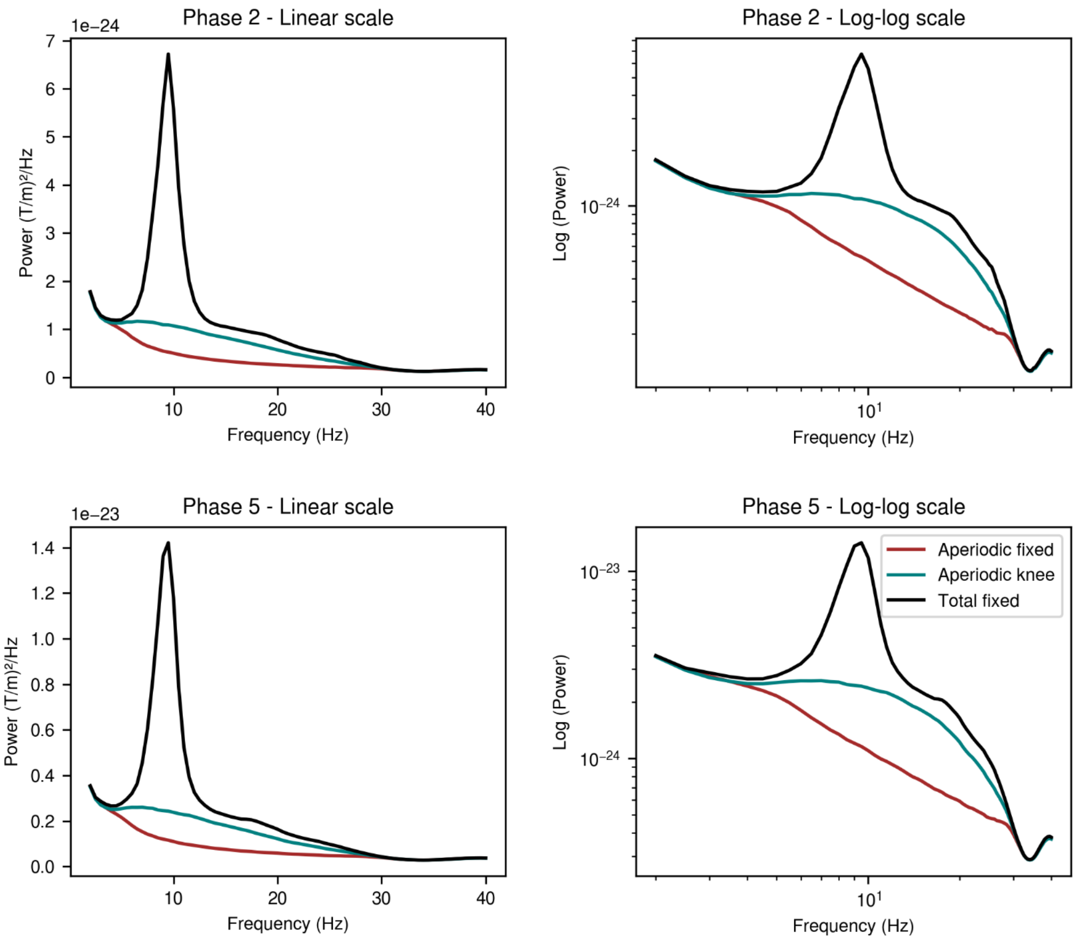


**Supplementary Figure 3. Comparison of MEG aperiodic power spectrum with and without the knee parameter.** The curve of the aperiodic spectrum with knee (teal) is closer to the total spectrum (black) than the fixed-mode aperiodic spectrum (brown), but this may be through also capturing the beta band peak.

Supplementary Figure 4. Comparison of MEG spectral parametrisation between fixed and knee aperiodic modes. The fixed mode showed a better goodness-of-fit (r squared). Additionally, the knee mode did not improve the detection of Theta peaks.

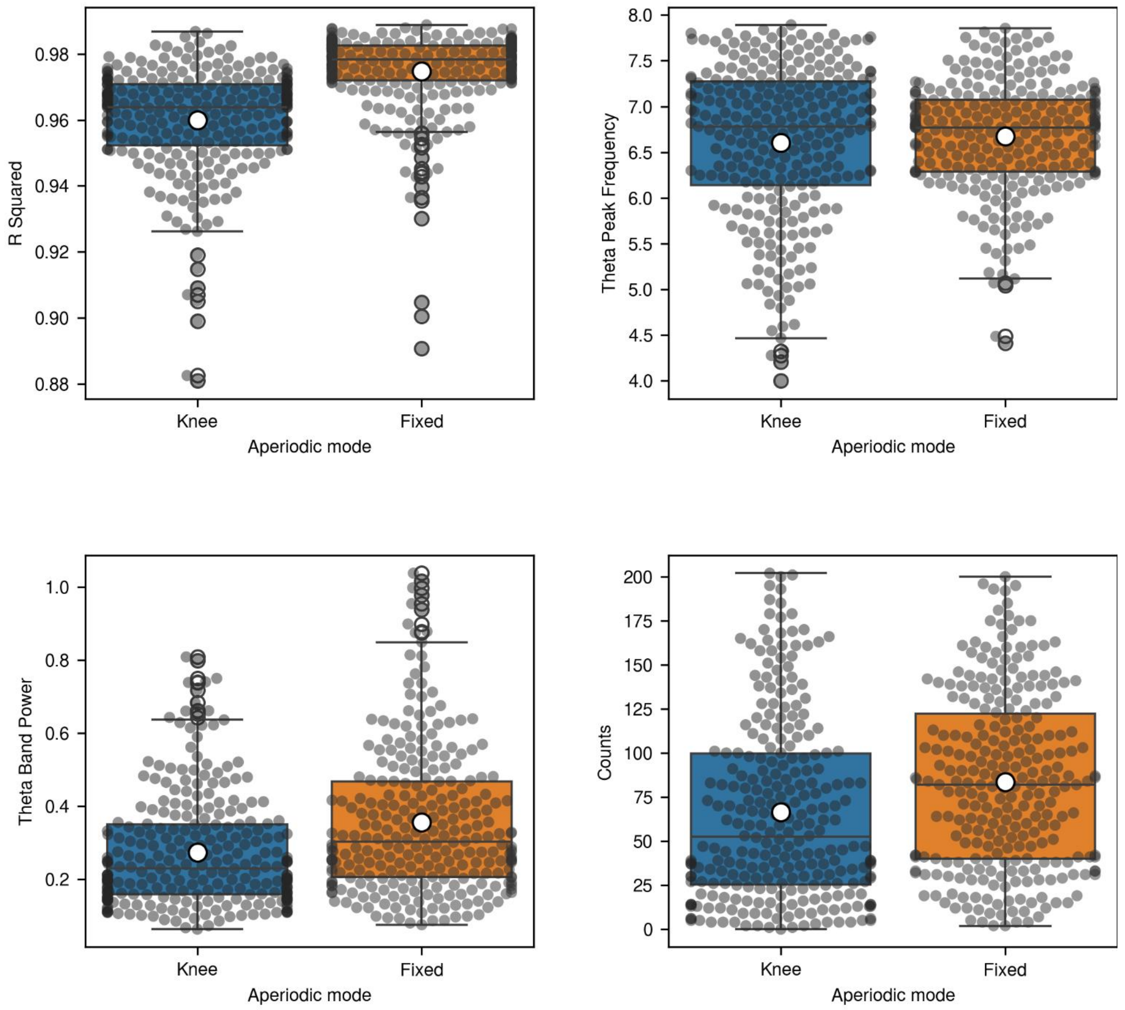


**Supplementary Figure 4. Comparison of MEG spectral parametrisation between fixed and knee aperiodic modes.** The fixed mode showed a better goodness-of-fit (r squared). Additionally, the knee mode did not improve the detection of Theta peaks.

Supplementary Figure 5. Comparison of the Aperiodic Exponent estimated using the fixed and knee aperiodic modes. The barplot of the Exponent with the knee mode (bottom left) does not show a flattening with age; instead, it shows an interaction effect, much like the Beta power estimated with the fixed mode (see Figure 4, bottom left). The topography of the Exponent with the knee mode (bottom right) also looks similar to the topography of Beta power estimated with the fixed mode (see Figure 4, bottom middle), rather than the more typical widespread distribution**.**

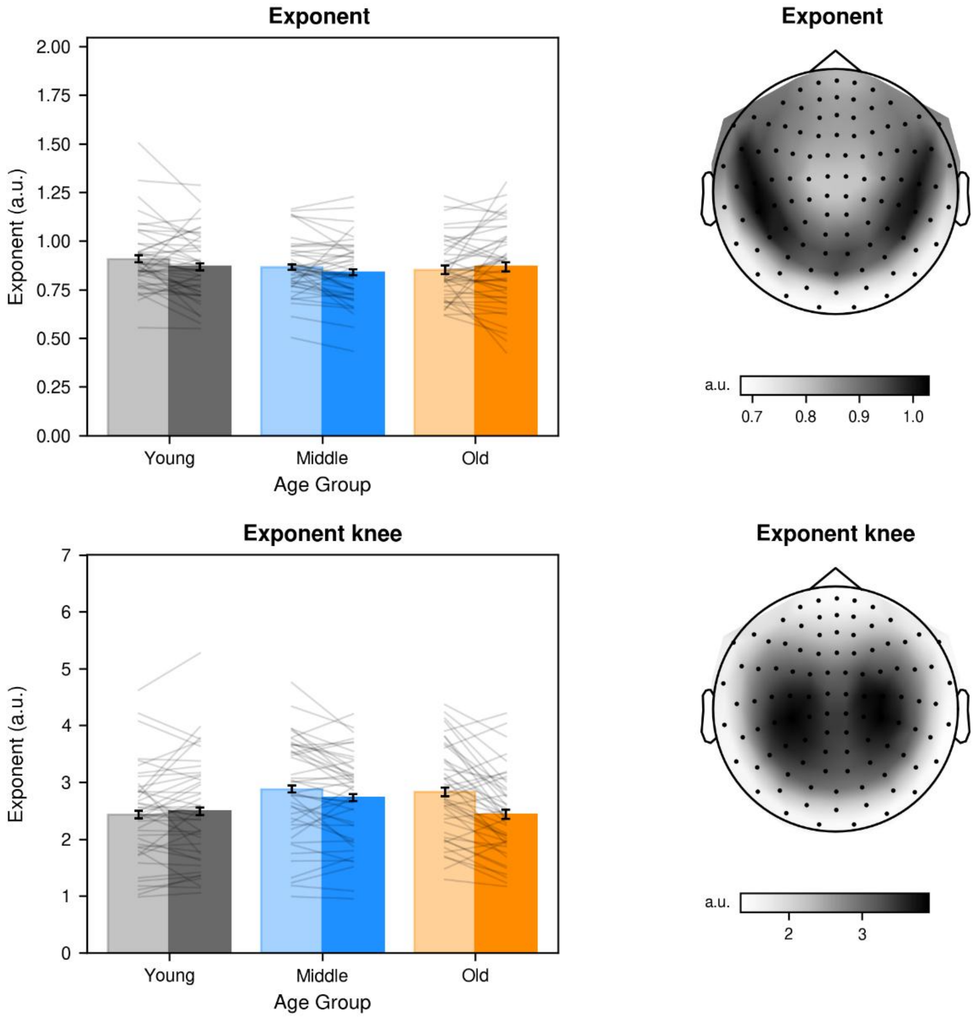


**Supplementary Figure 5. Comparison of the Aperiodic Exponent estimated using the fixed and knee aperiodic modes.** The barplot of the Exponent with the knee mode (bottom left) does not show a flattening with age; instead, it shows an interaction effect, much like the Beta power estimated with the fixed mode (see Figure 4, bottom left). The topography of the Exponent with the knee mode (bottom right) also looks similar to the topography of Beta power estimated with the fixed mode (see Figure 4, bottom middle), rather than the more typical widespread distribution.

Supplementary Figure 6. Comparison of ECG aperiodic power spectrum with and without the knee parameter. The aperiodic spectrum with knee (teal) is closer to the total spectrum (black) than the fixed-mode aperiodic spectrum (brown), and in this case (unlike MEG above), there should be no influence of brain oscillations like Beta.

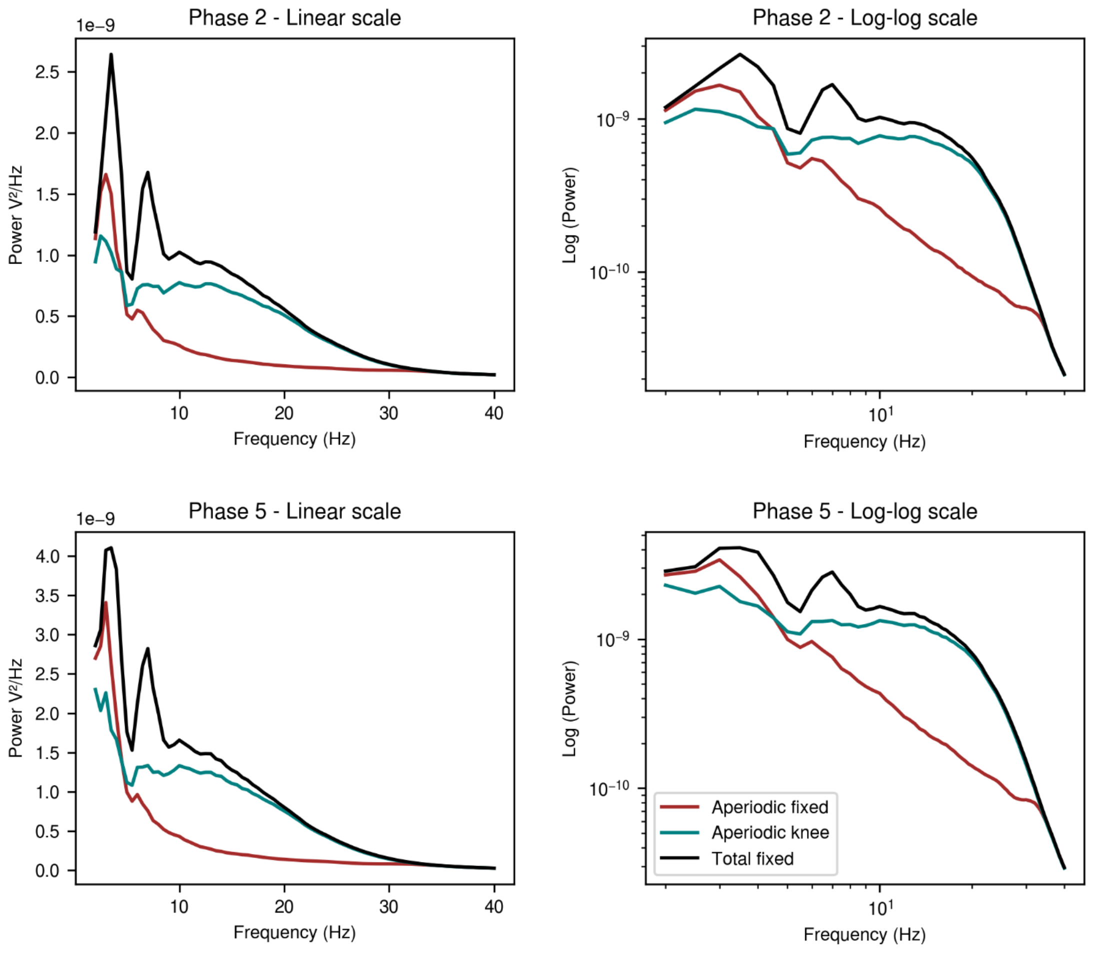


**Supplementary Figure 6. Comparison of ECG aperiodic power spectrum with and without the knee parameter.** The aperiodic spectrum with knee (teal) is closer to the total spectrum (black) than the fixed-mode aperiodic spectrum (brown), and in this case (unlike MEG above), there should be no influence of brain oscillations like Beta.

#### Supplementary Results

Supplementary Table 1: Low-Alpha (8-10 Hz) band (gradiometers) statistics for the three fixed effects of baseline age (A0), change in age (dA) and their interaction (A0:dA) from the four Linear Mixed Effects (LME) models. Abbreviations: a.u. = arbitrary units, p.a. = per annum (for main effects of A0 and dA).

| | Low-Alpha Power | | | Low-Alpha Frequency | | |
|---|---|---|---|---|---|---|
| **Model / Age Effect** | **Parameter Estimate, a.u. p.a. (SD)** | **T-statistic (df)** | **P-value uncorrected (corrected)** | **Parameter Estimate, Hz p.a. (SD)** | **T-statistic (df)** | **P-value uncorrected (corrected)** |
| **Basic Model** | | | | | | |
| **A0** | 0.0013 (0.0018) | 0.75 (131.1) | 0.45 (1) | -0.0055 (0.002) | -2.8 (131.4) | 0.0059 (0.069) |
| **dA** | -0.0022 (0.0013) | -1.68 (131.5) | 0.096 (0.73) | -0.0041 (0.0029) | -1.4 (132.9) | 0.17 (0.89) |
| **A0:dA** | -0.00024 ($9.4 \times 10^{-5}$) | -2.54 (131.7) | 0.012 (0.17) | -0.00037 (0.00021) | -1.79 (133.7) | 0.075 (0.64) |
| **Empty-room Model** | | | | | | |
| **A0** | 0.004 (0.0057) | 0.71 (124.0) | 0.48 (1) | -0.0055 (0.002) | -2.8 (131.4) | 0.0059 (0.069) |
| **dA** | -0.0079 (0.0044) | -1.77 (124.8) | 0.078 (0.66) | -0.0041 (0.0029) | -1.4 (132.9) | 0.17 (0.9) |
| **A0:dA** | -0.00081 (0.00031) | -2.6 (124.5) | 0.01 (0.14) | -0.00037 (0.00021) | -1.79 (133.7) | 0.075 (0.65) |
| **Cardiac Model** | | | | | | |
| **A0** | 0.0051 (0.0058) | 0.89 (133.7) | 0.38 (1) | -0.0055 (0.002) | -2.8 (131.4) | 0.0059 (0.07) |
| **dA** | -0.0072 (0.0043) | -1.67 (130.9) | 0.097 (0.74) | -0.0041 (0.0029) | -1.4 (132.9) | 0.17 (0.89) |
| **A0:dA** | -0.00078 (0.00031) | -2.55 (131.1) | 0.012 (0.16) | -0.00037 (0.00021) | -1.79 (133.7) | 0.075 (0.65) |
| **6-covariate Model** | | | | | | |
| **A0** | 0.0014 (0.0017) | 0.81 (129.9) | 0.42 (1) | -0.005 (0.002) | -2.49 (131.0) | 0.014 (0.16) |
| **dA** | -0.0023 (0.0015) | -1.56 (132.2) | 0.12 (0.8) | -0.0033 (0.0033) | -1.02 (141.7) | 0.31 (0.99) |
| **A0:dA** | -0.00024 ($9.4 \times 10^{-5}$) | -2.52 (127.6) | 0.013 (0.17) | -0.00038 (0.00021) | -1.83 (129.9) | 0.069 (0.61) |

Supplementary Table 2: High-Alpha (10-12 Hz) band (gradiometer) statistics for the three fixed effects of baseline age (A0), change in age (dA) and their interaction (A0:dA) from the four Linear Mixed Effects (LME) models. Abbreviations: a.u. = arbitrary units, p.a. = per annum (for main effects of A0 and dA). Effects that survived correction in bold.

| | High-Alpha Power | | | High-Alpha Frequency | | |
|---|---|---|---|---|---|---|
| **Model / Age Effect** | **Parameter Estimate, a.u. p.a. (SD)** | **T-statistic (df)** | **P-value uncorrected (corrected)** | **Parameter Estimate, Hz p.a. (SD)** | **T-statistic (df)** | **P-value uncorrected (corrected)** |
| **Basic Model** | | | | | | |
| **A0** | -0.0035 (0.0013) | -2.73 (131.1) | 0.0072 (0.084) | 0.0043 (0.0017) | 2.5 (131.3) | 0.014 (0.16) |
| **dA** | -0.0057 (0.0011) | -4.99 (131.7) | $1.9 \times 10^{-6}$ **($< 10^{-4}$)** | 0.0023 (0.0023) | 1.02 (132.5) | 0.31 (0.99) |
| **A0:dA** | $-5.4 \times 10^{-6}$ ($8.1 \times 10^{-5}$) | -0.07 (132.0) | 0.95 (1) | $9.9 \times 10^{-5}$ (0.00016) | 0.62 (133.1) | 0.54 (1) |
| **Empty-room Model** | | | | | | |
| **A0** | -0.017 (0.0055) | -3.03 (133.5) | 0.0029 **(0.034)** | 0.0043 (0.0017) | 2.5 (131.3) | 0.014 (0.16) |
| **dA** | -0.026 (0.0049) | -5.27 (131.7) | $5.3 \times 10^{-7}$ **($< 10^{-4}$)** | 0.0023 (0.0023) | 1.02 (132.5) | 0.31 (0.99) |
| **A0:dA** | -0.00017 (0.00035) | -0.48 (133.1) | 0.63 (1) | $9.9 \times 10^{-5}$ (0.00016) | 0.62 (133.1) | 0.54 (1) |
| **Cardiac Model** | | | | | | |
| **A0** | -0.015 (0.0055) | -2.68 (131.5) | 0.0084 (0.1) | 0.0043 (0.0017) | 2.5 (131.3) | 0.014 (0.16) |
| **dA** | -0.024 (0.005) | -4.87 (130.6) | $3.1 \times 10^{-6}$ **($< 10^{-4}$)** | 0.0023 (0.0023) | 1.02 (132.5) | 0.31 (0.99) |
| **A0:dA** | $-7.9 \times 10^{-6}$ (0.00035) | -0.02 (130.7) | 0.98 (1) | $9.9 \times 10^{-5}$ (0.00016) | 0.62 (133.1) | 0.54 (1) |
| **6-covariate Model** | | | | | | |
| **A0** | -0.003 (0.0013) | -2.36 (130.7) | 0.02 (0.21) | 0.0044 (0.0018) | 2.51 (131.2) | 0.013 (0.15) |
| **dA** | -0.0076 (0.0012) | -6.18 (133.9) | $7.2 \times 10^{-9}$ **($< 10^{-4}$)** | 0.0034 (0.0025) | 1.39 (139.2) | 0.17 (0.9) |
| **A0:dA** | $-1.5 \times 10^{-5}$ ($7.7 \times 10^{-5}$) | -0.2 (128.5) | 0.85 (1) | 0.00013 (0.00016) | 0.8 (129.4) | 0.43 (1) |

Supplementary Table 3: Low-Beta (12-20 Hz) band (gradiometer) statistics for the three fixed effects of baseline age (A0), change in age (dA) and their interaction (A0:dA) from the four Linear Mixed Effects (LME) models. Abbreviations: a.u. = arbitrary units, p.a. = per annum (for main effects of A0 and dA). Effects that survived correction in bold.

| | **Low-Beta Power** | | | **Low-Beta Frequency** | | |
|---|---|---|---|---|---|---|
| **Model / Age Effect** | **Parameter Estimate, a.u. p.a. (SD)** | **T-statistic (df)** | **P-value uncorrected (corrected)** | **Parameter Estimate, Hz p.a. (SD)** | **T-statistic (df)** | **P-value uncorrected (corrected)** |
| **Basic Model** | | | | | | |
| **A0** | 0.0018 (0.0008) | 2.21 (131.1) | 0.029 (0.3) | -0.0017 (0.0057) | -0.29 (131.2) | 0.77 (1) |
| **dA** | -0.0019 (0.00059) | -3.22 (131.4) | 0.0016 **(0.026)** | -0.018 (0.0068) | -2.57 (132.2) | 0.011 (0.16) |
| **A0:dA** | -0.00013 ($4.2 \times 10^{-5}$) | -3.21 (131.6) | 0.0016 **(0.022)** | -0.0011 (0.00048) | -2.3 (132.7) | 0.023 (0.28) |
| **Empty-room Model** | | | | | | |
| **A0** | 0.013 (0.0057) | 2.22 (129.2) | 0.028 (0.29) | -0.0017 (0.0057) | -0.29 (131.2) | 0.77 (1) |
| **dA** | -0.013 (0.0042) | -3.1 (130.1) | 0.0024 **(0.039)** | -0.018 (0.0068) | -2.57 (132.2) | 0.011 (0.16) |
| **A0:dA** | -0.00093 (0.0003) | -3.13 (129.4) | 0.0021 **(0.03)** | -0.0011 (0.00048) | -2.3 (132.7) | 0.023 (0.28) |
| **Cardiac Model** | | | | | | |
| **A0** | 0.011 (0.0058) | 2.0 (130.3) | 0.048 (0.44) | -0.0017 (0.0057) | -0.29 (131.2) | 0.77 (1) |
| **dA** | -0.015 (0.0041) | -3.72 (129.9) | 0.00029 **(0.0044)** | -0.018 (0.0068) | -2.57 (132.2) | 0.011 (0.16) |
| **A0:dA** | -0.001 (0.00029) | -3.54 (129.0) | 0.00055 **(0.0077)** | -0.0011 (0.00048) | -2.3 (132.7) | 0.023 (0.28) |
| **6-covariate Model** | | | | | | |
| **A0** | 0.0019 (0.0008) | 2.43 (130.0) | 0.017 (0.18) | -0.002 (0.0057) | -0.35 (129.8) | 0.73 (1) |
| **dA** | -0.0023 (0.00066) | -3.52 (131.8) | 0.0006 **(0.0089)** | -0.01 (0.0077) | -1.35 (137.2) | 0.18 (0.92) |
| **A0:dA** | -0.00014 ($4.1 \times 10^{-5}$) | -3.4 (127.7) | 0.00089 **(0.013)** | -0.0012 (0.00048) | -2.42 (128.0) | 0.017 (0.22) |

Supplementary Table 4: High-Beta (20-30 Hz) band (gradiometer) statistics for the three fixed effects of baseline age (A0), change in age (dA) and their interaction (A0:dA) from the four Linear Mixed Effects (LME) models. Abbreviations: a.u. = arbitrary units, p.a. = per annum (for main effects of A0 and dA). Effects that survived correction in bold.

| | **High-Beta Power** | | | **High-Beta Frequency** | | |
|---|---|---|---|---|---|---|
| **Model / Age Effect** | **Parameter Estimate, a.u. p.a. (SD)** | **T-statistic (df)** | **P-value uncorrected (corrected)** | **Parameter Estimate, Hz p.a. (SD)** | **T-statistic (df)** | **P-value uncorrected (corrected)** |
| **Basic Model** | | | | | | |
| **A0** | 0.0005 (0.00059) | 0.84 (131.1) | 0.4 (1) | -0.011 (0.0055) | -1.99 (131.2) | 0.049 (0.45) |
| **dA** | -0.0035 (0.00049) | -7.15 (131.6) | $5.3 \times 10^{-11}$ **($< 10^{-4}$)** | -0.017 (0.0052) | -3.31 (131.8) | 0.0012 **(0.019)** |
| **A0:dA** | -0.00023 ($3.5 \times 10^{-5}$) | -6.7 (131.8) | $5.6 \times 10^{-10}$ **($< 10^{-4}$)** | $9.7 \times 10^{-5}$ (0.00037) | 0.26 (132.1) | 0.79 (1) |
| **Empty-room Model** | | | | | | |
| **A0** | 0.0045 (0.0056) | 0.81 (130.2) | 0.42 (1) | -0.011 (0.0055) | -1.99 (131.2) | 0.049 (0.45) |
| **dA** | -0.034 (0.0058) | -5.84 (157.4) | $2.9 \times 10^{-8}$ **($< 10^{-4}$)** | -0.017 (0.0052) | -3.31 (131.8) | 0.0012 **(0.019)** |
| **A0:dA** | -0.0022 (0.00033) | -6.61 (130.3) | $9 \times 10^{-10}$ **($< 10^{-4}$)** | $9.7 \times 10^{-5}$ (0.00037) | 0.26 (132.1) | 0.79 (1) |
| **Cardiac Model** | | | | | | |
| **A0** | 0.0057 (0.0055) | 1.05 (132.3) | 0.3 (0.98) | -0.011 (0.0055) | -1.99 (131.2) | 0.049 (0.45) |
| **dA** | -0.03 (0.005) | -5.94 (139.6) | $2.2 \times 10^{-8}$ **($< 10^{-4}$)** | -0.017 (0.0052) | -3.31 (131.8) | 0.0012 **(0.019)** |
| **A0:dA** | -0.0021 (0.00033) | -6.19 (132.6) | $7.2 \times 10^{-9}$ **($< 10^{-4}$)** | $9.7 \times 10^{-5}$ (0.00037) | 0.26 (132.1) | 0.79 (1) |
| **6-covariate Model** | | | | | | |
| **A0** | 0.00074 (0.00059) | 1.26 (130.2) | 0.21 (0.93) | -0.012 (0.0056) | -2.18 (131.1) | 0.031 (0.31) |
| **dA** | -0.0037 (0.00056) | -6.65 (133.3) | $6.8 \times 10^{-10}$ **($< 10^{-4}$)** | -0.017 (0.0059) | -2.94 (135.2) | 0.0039 (0.058) |
| **A0:dA** | -0.00024 ($3.5 \times 10^{-5}$) | -6.76 (128.0) | $4.3 \times 10^{-10}$ **($< 10^{-4}$)** | $9.7 \times 10^{-5}$ (0.00037) | 0.26 (128.9) | 0.79 (1) |

Supplementary Figure 7. Topographic scalp distributions of Alpha and Beta sub-bands power. Periodic peaks detected with the spectral parametrisation algorithm were assigned to an Alpha or Beta sub-band, based on their centre frequency (see Methods). To obtain a sub-band topography, peaks' power parameters were averaged across all datasets (subjects and sessions) for each MEG channel. Datasets without detected peaks in the sub-band were excluded from the average. Low- and High-Alpha power distributions look identical, which does not support a special dissociation of the Alpha sub-bands, despite the differences in age-related effects. High-Beta peak power is more anterior than Low-Beta peaks, consistent with sources in dorsal premotor and posterior motor areas, respectively (Nougaret et al., 2024).

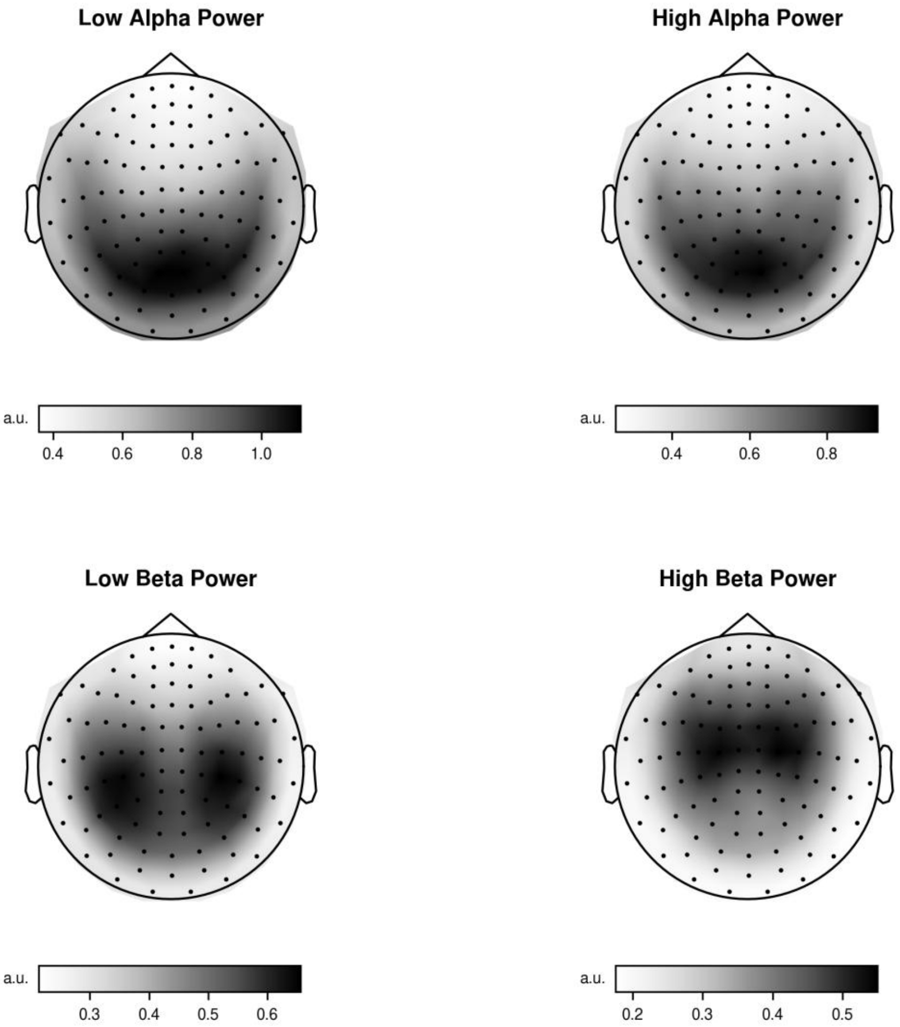


**Supplementary Figure 7. Topographic scalp distributions of Alpha and Beta sub-bands power.** Periodic peaks detected with the spectral parametrisation algorithm were assigned to an Alpha or Beta sub-band, based on their center frequency (see Methods). To obtain a sub-band topography, peaks' power parameters were averaged across all datasets (subjects and sessions) for each MEG channel. Datasets without detected peaks in the sub-band were excluded from the average. Low- and High-Alpha power distributions look identical, which does not support a special dissociation of the Alpha sub-bands, despite the differences in age-related effects. High-Beta peak power is more anterior than Low-Beta peaks, consistent with sources in dorsal premotor and posterior motor areas, respectively (Nougaret et al., 2024).

Supplementary Table 5. Effects of age on empty-room spectral features (gradiometers, fixed mode). All parameters statistics for the three fixed effects of baseline age (A0), change in age (dA) and their interaction (A0:dA) from the four Linear Mixed Effects (LME) models. Abbreviations: a.u. = arbitrary units (for exponent or power estimates), p.a. = per annum (for main effects of A0 and dA). Effects that survived correction in bold.

| Parameter / Age effect | Parameter Estimate, a.u./Hz p.a. (SD) | T-statistic (df) | P-value uncorrected (corrected) |
|---|---|---|---|
| **Exponent** | | | |
| **A0** | 0.00037 (0.00015) | 2.39 (132.1) | 0.018 (0.18) |
| **dA** | 0.00096 (0.0004) | 2.4 (136.6) | 0.018 (0.21) |
| **A0:dA** | -1.6 x $10^{-5}$ (2.8 x $10^{-5}$) | -0.56 (139.1) | 0.57 (1) |
| **Theta Peak Frequency** | | | |
| **A0** | 0.0012 (0.0012) | 1.01 (262.0) | 0.32 (0.99) |
| **dA** | 0.0016 (0.0032) | 0.5 (262.0) | 0.62 (1) |
| **A0:dA** | 2.7 x $10^{-5}$ (0.00023) | 0.12 (262.0) | 0.91 (1) |
| **Theta Band Power** | | | |
| **A0** | -2.2 x $10^{-5}$ (8.1 x $10^{-5}$) | -0.27 (132.2) | 0.79 (1) |
| **dA** | -0.00097 (0.00021) | -4.6 (136.8) | 9.5 x $10^{-6}$ **(0.0001)** |
| **A0:dA** | 8.6 x 10-7 (1.5 x $10^{-5}$) | 0.06 (139.3) | 0.95 (1) |
| **Alpha Peak Frequency** | | | |
| **A0** | -0.0013 (0.0015) | -0.85 (132.0) | 0.4 (1) |
| **dA** | 0.0093 (0.0038) | 2.48 (136.0) | 0.014 (0.18) |
| **A0:dA** | 0.00027 (0.00026) | 1.03 (138.2) | 0.31 (0.99) |
| **Alpha Band Power** | | | |
| **A0** | -0.00013 (0.00019) | -0.66 (131.7) | 0.51 (1) |
| **dA** | 0.0021 (0.00039) | 5.39 (134.4) | 3.1 x 10-7 **(< $10^{-4}$)** |
| **A0:dA** | -5.9 x $10^{-6}$ (2.7 x $10^{-5}$) | -0.22 (136.0) | 0.83 (1) |
| **Beta Peak Frequency** | | | |
| **A0** | 0.006 (0.0067) | 0.88 (131.0) | 0.38 (1) |
| **dA** | -0.29 (0.018) | -16.51 (135.7) | 2.9 x 10-34 **(< $10^{-4}$)** |
| **A0:dA** | 3.5 x $10^{-5}$ (0.0013) | 0.03 (138.3) | 0.98 (1) |
| **Beta Band Power** | | | |
| **A0** | 0.0002 (0.00023) | 0.85 (131.9) | 0.39 (1) |
| **dA** | -0.0034 (0.0006) | -5.63 (136.3) | 9.8 x 10-8 **(< $10^{-4}$)** |
| **A0:dA** | -7 x $10^{-5}$ (4.2 x $10^{-5}$) | -1.65 (138.8) | 0.1 (0.73) |
| **Low Alpha Peak Frequency** | | | |
| **A0** | 0.00031 (0.00088) | 0.35 (132.1) | 0.73 (1) |
| **dA** | 0.0036 (0.0022) | 1.67 (136.2) | 0.098 (0.73) |
| **A0:dA** | 0.00015 (0.00015) | 1.01 (138.5) | 0.31 (0.99) |
| **Low Alpha Band Power** | | | |
| **A0** | -7.5 x $10^{-5}$ (0.0002) | -0.38 (131.6) | 0.71 (1) |
| **dA** | 0.0015 (0.00039) | 3.75 (134.2) | 0.00027 **(0.0042)** |
| **A0:dA** | 2.1 x $10^{-6}$ (2.7 x $10^{-5}$) | 0.08 (135.7) | 0.94 (1) |
| **High Alpha Peak Frequency** | | | |
| **A0** | 9.3 x $10^{-5}$ (0.00094) | 0.1 (262.0) | 0.92 (1) |
| **dA** | 0.0037 (0.0025) | 1.47 (262.0) | 0.14 (0.86) |
| **A0:dA** | 5.6 x $10^{-5}$ (0.00018) | 0.32 (262.0) | 0.75 (1) |
| **High Alpha Band Power** | | | |
| **A0** | -0.00018 (0.00018) | -0.98 (131.8) | 0.33 (0.99) |
| **dA** | 0.0023 (0.00039) | 5.84 (134.8) | 3.8 x 10-8 **(< $10^{-4}$)** |
| **A0:dA** | -3.2 x $10^{-6}$ (2.8 x $10^{-5}$) | -0.12 (136.5) | 0.91 (1) |
| **Low Beta Peak Frequency** | | | |
| **A0** | 0.00045 (0.0022) | 0.21 (131.9) | 0.84 (1) |
| **dA** | -0.054 (0.0056) | -9.6 (136.2) | 5.4 x 10-17 **(< $10^{-4}$)** |
| **A0:dA** | 0.00037 (0.00039) | 0.95 (138.6) | 0.34 (0.99) |
| **Low Beta Band Power** | | | |
| **A0** | -4.6 x $10^{-5}$ (0.00021) | -0.22 (131.8) | 0.83 (1) |
| **dA** | 0.0028 (0.00048) | 5.87 (135.4) | 3.2 x 10-8 **(< $10^{-4}$)** |
| **A0:dA** | -4.5 x $10^{-6}$ (3.4 x $10^{-5}$) | -0.13 (137.4) | 0.89 (1) |
| **High Beta Peak Frequency** | | | |
| **A0** | 0.0013 (0.0024) | 0.55 (130.5) | 0.58 (1) |
| **dA** | -0.022 (0.0065) | -3.44 (135.2) | 0.00077 **(0.011)** |
| **A0:dA** | 0.0002 (0.00045) | 0.43 (137.9) | 0.67 (1) |
| **High Beta Band Power** | | | |
| **A0** | 0.00023 (0.00019) | 1.22 (131.3) | 0.23 (0.94) |

| | | | |
|---|---|---|---|
| **dA** | -0.0069 (0.00049) | -14.15 (135.8) | 1.7 x 10-28 **(< $10^{-4}$)** |
| **A0:dA** | -7.7 x $10^{-5}$ (3.4 x $10^{-5}$) | -2.24 (138.4) | 0.027 (0.29) |
| **Gamma Peak Frequency** | | | |
| **A0** | 0.0055 (0.0019) | 2.84 (262.0) | 0.0048 (0.057) |
| **dA** | 0.045 (0.0051) | 8.74 (262.0) | 2.8 x 10-16 **(< $10^{-4}$)** |
| **A0:dA** | -9.6 x $10^{-5}$ (0.00036) | -0.27 (262.0) | 0.79 (1) |
| **Gamma Band Power** | | | |
| **A0** | -0.00015 (0.00012) | -1.26 (262.0) | 0.21 (0.93) |
| **dA** | -0.0026 (0.00032) | -8.34 (262.0) | 4.2 x 10-15 **(< $10^{-4}$)** |
| **A0:dA** | 1.7 x $10^{-5}$ (2.2 x $10^{-5}$) | 0.75 (262.0) | 0.45 (1) |

Supplementary Figure 8. Differences in empty-room power spectrum between experimental sessions. The colour lines represent the mean empty-room power spectrum across datasets for each phase. The colour areas above and below the lines represent the standard error of the mean for the between-phase differences. Power differences are likely to be due mainly to the upgrade of the MEG system (see more details in Methods) but could also owe to changes in environmental noise over the years intervening between Phases 2 and 5. After the system change, the noise power decreased in Theta, High-Beta and Gamma frequency ranges, but increased in Alpha and Low-Beta frequency ranges.

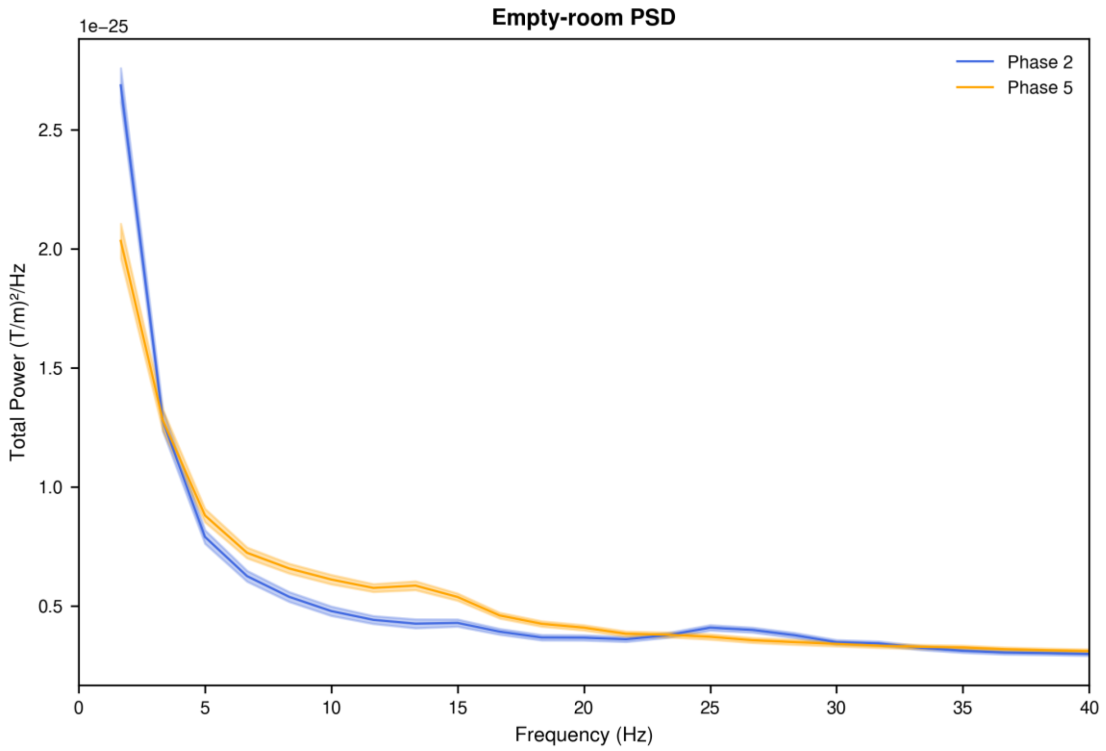


**Supplementary Figure 8. Differences in empty-room power spectrum between experimental sessions.** The colour lines represent the mean empty-room power spectrum across datasets for each phase. The colour areas above and below the lines represent the standard error of the mean for the between-phase differences. Power differences are likely to be due mainly to the upgrade of the MEG system (see more details in Methods) but could also owe to changes in environmental noise over the years intervening between Phases 2 and 5. After the system change, the noise power decreased in Theta, High-Beta and Gamma frequency ranges, but increased in Alpha and Low-Beta frequency ranges.

Supplementary Table 6. Effects of age on ECG spectral parameters. All parameters statistics for the three fixed effects of baseline age (A0), change in age (dA) and their interaction (A0:dA) from the four Linear Mixed Effects (LME) models. Several parameters are missing from the table because there were not enough detections across datasets, and therefore, they could not be tested. Abbreviations: a.u. = arbitrary units (for exponent or power estimates), p.a. = per annum (for main effects of A0 and dA). Effect that survived correction in bold.

| Parameter / Age effect | Parameter Estimate, a.u./Hz p.a. (SD) | T-statistic (df) | P-value uncorrected (corrected) |
|---|---|---|---|
| **Exponent** | | | |
| **A0** | -0.031 (0.011) | -2.91 (131.3) | 0.0043 **(0.0087)** |
| **dA** | -0.018 (0.013) | -1.41 (132.3) | 0.16 (0.41) |
| **A0:dA** | -0.00096 (0.00092) | -1.04 (132.9) | 0.3 (0.66) |
| **Theta Peak Frequency** | | | |
| **A0** | -0.001 (0.0055) | -0.19 (129.0) | 0.85 (1) |
| **dA** | -0.0057 (0.0086) | -0.67 (128.6) | 0.51 (0.88) |
| **A0:dA** | -0.0011 (0.0006) | -1.8 (128.3) | 0.074 (0.21) |
| **Theta Band Power** | | | |
| **A0** | -0.00025 (0.00068) | -0.38 (127.0) | 0.71 (0.97) |
| **dA** | -0.00026 (0.0011) | -0.24 (126.8) | 0.81 (0.99) |
| **A0:dA** | $-5.8 \times 10^{-5}$ ($7.5 \times 10^{-5}$) | -0.78 (126.4) | 0.44 (0.82) |

Supplementary Table 7. Effects of age on MEG cardiac components (gradiometers, fixed mode). All parameters statistics for the three fixed effects of baseline age (A0), change in age (dA) and their interaction (A0:dA) from the four Linear Mixed Effects (LME) models. Abbreviations: a.u. = arbitrary units (for exponent or power estimates), p.a. = per annum (for main effects of A0 and dA). Effects that survived correction in bold.

| Parameter / Age effect | Parameter Estimate, a.u./Hz p.a. (SD) | T-statistic (df) | P-value uncorrected (corrected) |
|---|---|---|---|
| **Exponent** | | | |
| **A0** | 0.0015 (0.0014) | 1.08 (119.3) | 0.28 (0.93) |
| **dA** | -0.0026 (0.002) | -1.32 (102.0) | 0.19 (0.83) |
| **A0:dA** | -4.4 x $10^{-5}$ (0.00014) | -0.31 (103.4) | 0.75 (1) |
| **Theta Peak Frequency** | | | |
| **A0** | 0.0015 (0.0056) | 0.27 (126.6) | 0.79 (1) |
| **dA** | 0.012 (0.0089) | 1.36 (108.1) | 0.18 (0.8) |
| **A0:dA** | 0.00069 (0.00064) | 1.09 (110.9) | 0.28 (0.93) |
| **Theta Band Power** | | | |
| **A0** | 5.3 x $10^{-5}$ (0.00084) | 0.06 (118.3) | 0.95 (1) |
| **dA** | -0.0023 (0.0013) | -1.85 (98.1) | 0.067 (0.45) |
| **A0:dA** | -0.00023 (9.1 x $10^{-5}$) | -2.52 (100.6) | 0.013 (0.12) |
| **Alpha Peak Frequency** | | | |
| **A0** | 0.0037 (0.006) | 0.63 (108.2) | 0.53 (1) |
| **dA** | 0.0087 (0.013) | 0.68 (99.6) | 0.5 (1) |
| **A0:dA** | 0.00038 (0.0009) | 0.43 (103.0) | 0.67 (1) |
| **Alpha Band Power** | | | |
| **A0** | -0.0013 (0.00091) | -1.46 (108.8) | 0.15 (0.7) |
| **dA** | -0.0023 (0.0013) | -1.87 (81.8) | 0.066 (0.44) |
| **A0:dA** | -0.00015 (8.9 x $10^{-5}$) | -1.69 (83.6) | 0.095 (0.57) |
| **Beta Peak Frequency** | | | |
| **A0** | 0.011 (0.02) | 0.58 (117.9) | 0.57 (1) |
| **dA** | -0.016 (0.04) | -0.39 (110.3) | 0.7 (1) |
| **A0:dA** | 0.0019 (0.0029) | 0.65 (112.8) | 0.51 (1) |
| **Beta Band Power** | | | |
| **A0** | -0.0026 (0.001) | -2.63 (122.9) | 0.0097 (0.072) |
| **dA** | -0.0032 (0.0014) | -2.3 (105.4) | 0.023 (0.2) |
| **A0:dA** | -0.00031 (9.8 x $10^{-5}$) | -3.12 (106.8) | 0.0024 **(0.023)** |
| **Low Beta Peak Frequency** | | | |
| **A0** | -0.0022 (0.01) | -0.21 (117.3) | 0.83 (1) |
| **dA** | -0.06 (0.025) | -2.38 (115.6) | 0.019 (0.17) |
| **A0:dA** | -1.4 x $10^{-5}$ (0.0018) | -0.01 (119.1) | 0.99 (1) |
| **Low Beta Band Power** | | | |
| **A0** | -0.0021 (0.001) | -2.05 (121.4) | 0.042 (0.28) |
| **dA** | -0.0031 (0.0014) | -2.32 (100.9) | 0.022 (0.19) |
| **A0:dA** | -0.00029 (9.6 x $10^{-5}$) | -2.98 (102.4) | 0.0036 **(0.034)** |
| **High Beta Peak Frequency** | | | |
| **A0** | -0.0037 (0.0092) | -0.4 (116.9) | 0.69 (1) |
| **dA** | -0.02 (0.022) | -0.89 (115.2) | 0.37 (0.98) |
| **A0:dA** | 8 x $10^{-5}$ (0.0016) | 0.05 (117.1) | 0.96 (1) |
| **High Beta Band Power** | | | |
| **A0** | -0.0021 (0.00087) | -2.41 (123.3) | 0.017 (0.13) |
| **dA** | -0.0027 (0.0012) | -2.24 (103.1) | 0.028 (0.23) |
| **A0:dA** | -0.00029 (8.4 x $10^{-5}$) | -3.39 (103.8) | 0.001 **(0.01)** |

Supplementary Table 8. Effects of age on covariates (head position and movement). All parameters statistics for the three fixed effects of baseline age (A0), change in age (dA) and their interaction (A0:dA) from the four Linear Mixed Effects (LME) models. Abbreviations: p.a. = per annum (for main effects of A0 and dA). P-values were corrected for multiple comparisons using Bonferroni across the four tests. Effects that survived correction in bold.

| Covariate / Age Effect | Parameter Estimate mm p.a. (SD) | T-statistic (df) | P-value uncorrected (corrected*) |
|---|---|---|---|
| **Head position (x)** | | | |
| **A0** | -0.014 (0.013) | -1.07 (131.9) | 0.29 (1.1) |
| **dA** | -0.055 (0.03) | -1.85 (135.6) | 0.066 (0.26) |
| **A0:dA** | -0.0015 (0.0021) | -0.7 (137.6) | 0.48 (1.9) |
| **Head position (y)** | | | |
| **A0** | -0.0079 (0.026) | -0.31 (132.1) | 0.76 (3) |
| **dA** | -0.33 (0.065) | -5.03 (136.4) | $1.5 \times 10^{-6}$ **($6 \times 10^{-6}$)** |
| **A0:dA** | 0.0025 (0.0046) | 0.55 (138.8) | 0.58 (2.3) |
| **Head position (z)** | | | |
| **A0** | -0.08 (0.031) | -2.58 (131.6) | 0.011 **(0.044)** |
| **dA** | 0.23 (0.066) | 3.54 (134.7) | 0.00055 **(0.0022)** |
| **A0:dA** | 0.00093 (0.0047) | 0.2 (136.5) | 0.84 (3.4) |
| **Head movement** | | | |
| **A0** | 0.003 (0.003) | 0.98 (131.9) | 0.33 (1.3) |
| **dA** | -0.00096 (0.007) | -0.14 (135.5) | 0.89 (3.6) |
| **A0:dA** | -0.00015 (0.00049) | -0.3 (137.5) | 0.77 (3.1) |

Supplementary Table 9. Effects of age but ignoring peaks within 21.9-23.9 Hz (fixed aperiodic mode, gradiometers). All parameters statistics for the three fixed effects of baseline age (A0), change in age (dA) and their interaction (A0:dA) from the four Linear Mixed Effects (LME) models. Abbreviations: a.u. = arbitrary units (for exponent or power estimates), p.a. = per annum (for main effects of A0 and dA). Effects that survived correction in bold.

| Parameter / Age effect | Parameter Estimate, a.u./Hz p.a. (SD) | T-statistic (df) | P-value uncorrected (corrected) |
|---|---|---|---|
| **Exponent** | | | |
| **A0** | -0.0011 (0.00091) | -1.24 (131.2) | 0.22 (0.93) |
| **dA** | -0.0011 (0.00098) | -1.15 (131.9) | 0.25 (0.96) |
| **A0:dA** | 0.0002 (6.9 x $10^{-5}$) | 2.91 (132.4) | 0.0043 (0.055) |
| **Theta Peak Frequency** | | | |
| **A0** | 0.0031 (0.0031) | 0.99 (131.3) | 0.32 (0.99) |
| **dA** | -0.00097 (0.0048) | -0.2 (132.9) | 0.84 (1) |
| **A0:dA** | 0.00034 (0.00034) | 1.01 (133.8) | 0.32 (0.99) |
| **Theta Band Power** | | | |
| **A0** | 0.004 (0.0011) | 3.75 (131.2) | 0.00027 **(0.003)** |
| **dA** | 0.0033 (0.0011) | 3.1 (131.8) | 0.0024 **(0.034)** |
| **A0:dA** | 6.3 x $10^{-5}$ (7.5 x $10^{-5}$) | 0.84 (132.2) | 0.4 (1) |
| **Alpha Peak Frequency** | | | |
| **A0** | -0.016 (0.0034) | -4.64 (131.2) | 8.2 x $10^{-6}$ **(< $10^{-4}$)** |
| **dA** | -0.016 (0.0031) | -5.17 (131.7) | 8.5 x 10-7 **(< $10^{-4}$)** |
| **A0:dA** | -3.3 x $10^{-5}$ (0.00022) | -0.15 (132.0) | 0.88 (1) |
| **Alpha Band Power** | | | |
| **A0** | -0.0013 (0.0017) | -0.79 (131.1) | 0.43 (1) |
| **dA** | -0.006 (0.0012) | -5.15 (131.4) | 9.4 x 10-7 **(< $10^{-4}$)** |
| **A0:dA** | -0.00018 (8.3 x $10^{-5}$) | -2.12 (131.6) | 0.036 (0.37) |
| **Beta Peak Frequency** | | | |
| **A0** | -0.024 (0.013) | -1.88 (120.9) | 0.062 (0.49) |
| **dA** | -0.051 (0.01) | -5.13 (111.6) | 1.2 x $10^{-6}$ **(< $10^{-4}$)** |
| **A0:dA** | -0.00099 (0.00071) | -1.39 (110.7) | 0.17 (0.88) |
| **Beta Band Power** | | | |
| **A0** | 0.0017 (0.00081) | 2.08 (124.7) | 0.04 (0.35) |
| **dA** | -0.0033 (0.00063) | -5.21 (115.3) | 8.4 x 10-7 **(< $10^{-4}$)** |
| **A0:dA** | -0.0002 (4.5 x $10^{-5}$) | -4.38 (114.4) | 2.6 x $10^{-5}$ **(0.0002)** |
| **Low Alpha Peak Frequency** | | | |
| **A0** | -0.0055 (0.002) | -2.8 (131.4) | 0.0059 (0.064) |
| **dA** | -0.0041 (0.0029) | -1.4 (132.9) | 0.17 (0.88) |
| **A0:dA** | -0.00037 (0.00021) | -1.79 (133.7) | 0.075 (0.62) |
| **Low Alpha Band Power** | | | |
| **A0** | 0.0013 (0.0018) | 0.75 (131.1) | 0.45 (1) |
| **dA** | -0.0022 (0.0013) | -1.68 (131.5) | 0.096 (0.7) |
| **A0:dA** | -0.00024 (9.4 x $10^{-5}$) | -2.54 (131.7) | 0.012 (0.15) |
| **High Alpha Peak Frequency** | | | |
| **A0** | 0.0043 (0.0017) | 2.5 (131.3) | 0.014 (0.15) |
| **dA** | 0.0023 (0.0023) | 1.02 (132.5) | 0.31 (0.98) |
| **A0:dA** | 9.9 x $10^{-5}$ (0.00016) | 0.62 (133.1) | 0.54 (1) |
| **High Alpha Band Power** | | | |
| **A0** | -0.0035 (0.0013) | -2.73 (131.1) | 0.0072 (0.078) |
| **dA** | -0.0057 (0.0011) | -4.99 (131.7) | 1.9 x $10^{-6}$ **(< $10^{-4}$)** |
| **A0:dA** | -5.4 x $10^{-6}$ (8.1 x $10^{-5}$) | -0.07 (132.0) | 0.95 (1) |
| **Low Beta Peak Frequency** | | | |
| **A0** | -0.0017 (0.0057) | -0.29 (131.2) | 0.77 (1) |
| **dA** | -0.018 (0.0068) | -2.57 (132.2) | 0.011 (0.14) |
| **A0:dA** | -0.0011 (0.00048) | -2.3 (132.7) | 0.023 (0.26) |
| **Low Beta Band Power** | | | |
| **A0** | 0.0018 (0.0008) | 2.21 (131.1) | 0.029 (0.28) |
| **dA** | -0.0019 (0.00059) | -3.22 (131.4) | 0.0016 **(0.024)** |
| **A0:dA** | -0.00013 (4.2 x $10^{-5}$) | -3.21 (131.6) | 0.0016 **(0.02)** |
| **Gamma Peak Frequency** | | | |
| **A0** | -0.0036 (0.0063) | -0.56 (131.7) | 0.57 (1) |
| **dA** | 0.047 (0.015) | 3.22 (136.0) | 0.0016 **(0.024)** |
| **A0:dA** | 0.0014 (0.001) | 1.34 (137.2) | 0.18 (0.9) |
| **Gamma Band Power** | | | |
| **A0** | 7.6 x $10^{-5}$ (0.00013) | 0.61 (130.9) | 0.54 (1) |
| **dA** | -0.00058 (0.00025) | -2.34 (133.9) | 0.021 (0.24) |

| | | | |
|---|---|---|---|
| **A0:dA** | $1.1 \times 10^{-5}$ ($1.7 \times 10^{-5}$) | 0.65 (134.6) | 0.52 (1) |

Supplementary Table 10. Effects of age without power interpolation at 21.9-23.9 Hz (fixed aperiodic mode, gradiometers). All parameters statistics for the three fixed effects of baseline age (A0), change in age (dA) and their interaction (A0:dA) from the four Linear Mixed Effects (LME) models. Abbreviations: a.u. = arbitrary units (for exponent or power estimates), p.a. = per annum (for main effects of A0 and dA). Effects that survived correction in bold.

| Parameter / Age effect | Parameter Estimate, a.u./Hz p.a. (SD) | T-statistic (df) | P-value uncorrected (corrected) |
|---|---|---|---|
| **Exponent** | | | |
| **A0** | -0.0011 (0.00091) | -1.24 (131.2) | 0.22 (0.94) |
| **dA** | -0.0012 (0.00098) | -1.17 (131.9) | 0.24 (0.97) |
| **A0:dA** | 0.0002 (6.9 x $10^{-5}$) | 2.89 (132.4) | 0.0044 (0.064) |
| **Theta Peak Frequency** | | | |
| **A0** | 0.0036 (0.0031) | 1.14 (131.3) | 0.25 (0.97) |
| **dA** | -0.00072 (0.0047) | -0.15 (132.8) | 0.88 (1) |
| **A0:dA** | 0.00033 (0.00034) | 0.99 (133.7) | 0.32 (0.99) |
| **Theta Band Power** | | | |
| **A0** | 0.0041 (0.0011) | 3.79 (131.2) | 0.00023 **(0.0024)** |
| **dA** | 0.0036 (0.0011) | 3.32 (131.8) | 0.0012 **(0.02)** |
| **A0:dA** | 7.2 x $10^{-5}$ (7.6 x $10^{-5}$) | 0.94 (132.2) | 0.35 (0.99) |
| **Alpha Peak Frequency** | | | |
| **A0** | -0.016 (0.0034) | -4.54 (131.1) | 1.3 x $10^{-5}$ **(< $10^{-4}$)** |
| **dA** | -0.016 (0.0031) | -5.08 (131.7) | 1.3 x $10^{-6}$ **(0.0001)** |
| **A0:dA** | 2.4 x $10^{-5}$ (0.00022) | 0.11 (132.0) | 0.91 (1) |
| **Alpha Band Power** | | | |
| **A0** | -0.0013 (0.0017) | -0.8 (131.1) | 0.43 (1) |
| **dA** | -0.0061 (0.0012) | -5.17 (131.4) | 8.3 x 10-7 **(0.0001)** |
| **A0:dA** | -0.00018 (8.3 x $10^{-5}$) | -2.14 (131.6) | 0.035 (0.39) |
| **Beta Peak Frequency** | | | |
| **A0** | -0.029 (0.012) | -2.43 (131.1) | 0.016 (0.19) |
| **dA** | 0.011 (0.01) | 1.07 (131.5) | 0.29 (0.98) |
| **A0:dA** | -0.00068 (0.00071) | -0.95 (131.8) | 0.34 (0.99) |
| **Beta Band Power** | | | |
| **A0** | 0.0016 (0.00073) | 2.13 (131.1) | 0.035 (0.34) |
| **dA** | -0.0003 (0.00063) | -0.48 (131.6) | 0.63 (1) |
| **A0:dA** | -0.00021 (4.4 x $10^{-5}$) | -4.75 (131.9) | 5.2 x $10^{-6}$ **(0.0002)** |
| **Low Alpha Peak Frequency** | | | |
| **A0** | -0.0052 (0.002) | -2.66 (131.4) | 0.0089 (0.11) |
| **dA** | -0.0038 (0.0029) | -1.31 (132.9) | 0.19 (0.93) |
| **A0:dA** | -0.00034 (0.00021) | -1.66 (133.7) | 0.1 (0.75) |
| **Low Alpha Band Power** | | | |
| **A0** | 0.0013 (0.0017) | 0.77 (131.1) | 0.44 (1) |
| **dA** | -0.0021 (0.0013) | -1.55 (131.5) | 0.12 (0.82) |
| **A0:dA** | -0.00025 (9.4 x $10^{-5}$) | -2.61 (131.7) | 0.01 (0.14) |
| **High Alpha Peak Frequency** | | | |
| **A0** | 0.0044 (0.0017) | 2.57 (131.3) | 0.011 (0.13) |
| **dA** | 0.0015 (0.0023) | 0.64 (132.4) | 0.52 (1) |
| **A0:dA** | 0.00014 (0.00016) | 0.86 (133.1) | 0.39 (1) |
| **High Alpha Band Power** | | | |
| **A0** | -0.0036 (0.0013) | -2.8 (131.1) | 0.0059 (0.071) |
| **dA** | -0.0051 (0.0011) | -4.46 (131.7) | 1.8 x $10^{-5}$ **(0.0002)** |
| **A0:dA** | 4.7 x $10^{-6}$ (8 x $10^{-5}$) | 0.06 (132.0) | 0.95 (1) |
| **Low Beta Peak Frequency** | | | |
| **A0** | -0.0029 (0.0058) | -0.5 (131.2) | 0.62 (1) |
| **dA** | -0.014 (0.0068) | -2.04 (132.1) | 0.043 (0.45) |
| **A0:dA** | -0.0012 (0.00048) | -2.46 (132.6) | 0.015 (0.2) |
| **Low Beta Band Power** | | | |
| **A0** | 0.0018 (0.0008) | 2.25 (131.1) | 0.026 (0.28) |
| **dA** | -0.0017 (0.00059) | -2.92 (131.4) | 0.0041 (0.062) |
| **A0:dA** | -0.00014 (4.2 x $10^{-5}$) | -3.33 (131.6) | 0.0011 **(0.016)** |
| **High Beta Peak Frequency** | | | |
| **A0** | -0.0094 (0.0046) | -2.04 (131.2) | 0.044 (0.41) |
| **dA** | -0.033 (0.0054) | -6.1 (132.2) | 1.1 x 10-8 **(0.0001)** |
| **A0:dA** | 0.00023 (0.00038) | 0.6 (132.7) | 0.55 (1) |
| **High Beta Band Power** | | | |
| **A0** | 0.0006 (0.00058) | 1.03 (131.2) | 0.31 (0.99) |
| **dA** | 0.0013 (0.00064) | 2.02 (132.0) | 0.046 (0.47) |

| | | | |
|---|---|---|---|
| **A0:dA** | -0.00021 ($4.5 \times 10^{-5}$) | -4.59 (132.5) | $1 \times 10^{-5}$ **(0.0002)** |
| **Gamma Peak Frequency** | | | |
| **A0** | -0.0055 (0.0069) | -0.79 (133.2) | 0.43 (1) |
| **dA** | 0.041 (0.015) | 2.7 (135.1) | 0.0078 (0.11) |
| **A0:dA** | 0.0019 (0.0011) | 1.73 (138.0) | 0.086 (0.7) |
| **Gamma Band Power** | | | |
| **A0** | $8.1 \times 10^{-5}$ (0.00013) | 0.65 (132.1) | 0.52 (1) |
| **dA** | -0.00027 (0.00025) | -1.12 (133.2) | 0.27 (0.98) |
| **A0:dA** | $9.1 \times 10^{-6}$ ($1.7 \times 10^{-5}$) | 0.52 (135.7) | 0.6 (1) |

Supplementary Table 11. Effects of age from magnetometers (fixed aperiodic mode). Gradiometers only have higher SNR for superficial sources (since noise reduced by subtracting two coils), but have lower SNR for deep sources (since signal reduced by subtracting two coils) – relative to single coil magnetometers. All parameters statistics for the three fixed effects of baseline age (A0), change in age (dA) and their interaction (A0:dA) from the four Linear Mixed Effects (LME) models. Abbreviations: a.u. = arbitrary units (for exponent or power estimates), p.a. = per annum (for main effects of A0 and dA). Effects that survived correction in bold.

| Parameter / Age effect | Parameter Estimate, a.u./Hz p.a. (SD) | T-statistic (df) | P-value uncorrected (corrected) |
|---|---|---|---|
| **Exponent** | | | |
| **A0** | -0.0012 (0.001) | -1.17 (131.2) | 0.24 (0.96) |
| **dA** | -0.00038 (0.001) | -0.36 (131.9) | 0.72 (1) |
| **A0:dA** | 0.00022 (7.4 x $10^{-5}$) | 3.04 (132.3) | 0.0028 **(0.041)** |
| **Theta Peak Frequency** | | | |
| **A0** | 0.0042 (0.0033) | 1.25 (132.1) | 0.21 (0.94) |
| **dA** | 0.0011 (0.0059) | 0.19 (132.8) | 0.85 (1) |
| **A0:dA** | 0.00039 (0.00042) | 0.93 (133.6) | 0.36 (1) |
| **Theta Band Power** | | | |
| **A0** | 0.0045 (0.0012) | 3.84 (131.5) | 0.00019 **(0.0013)** |
| **dA** | 0.0039 (0.0014) | 2.71 (130.3) | 0.0076 (0.11) |
| **A0:dA** | 8.5 x $10^{-5}$ (0.0001) | 0.84 (130.6) | 0.4 (1) |
| **Alpha Peak Frequency** | | | |
| **A0** | -0.016 (0.0035) | -4.62 (131.2) | 9.1 x $10^{-6}$ **(< $10^{-4}$)** |
| **dA** | -0.019 (0.0038) | -4.95 (132.0) | 2.3 x $10^{-6}$ **(< $10^{-4}$)** |
| **A0:dA** | 5.8 x $10^{-5}$ (0.00027) | 0.22 (132.4) | 0.83 (1) |
| **Alpha Band Power** | | | |
| **A0** | -0.0018 (0.0018) | -1.02 (131.1) | 0.31 (0.99) |
| **dA** | -0.0065 (0.0013) | -5.14 (131.4) | 9.7 x 10-7 **(< $10^{-4}$)** |
| **A0:dA** | -0.00017 (9 x $10^{-5}$) | -1.88 (131.6) | 0.062 (0.58) |
| **Beta Peak Frequency** | | | |
| **A0** | -0.025 (0.014) | -1.77 (131.1) | 0.079 (0.62) |
| **dA** | -0.049 (0.011) | -4.38 (131.5) | 2.4 x $10^{-5}$ **(0.0004)** |
| **A0:dA** | -0.0013 (0.00079) | -1.7 (131.7) | 0.091 (0.72) |
| **Beta Band Power** | | | |
| **A0** | 0.0016 (0.00082) | 1.94 (131.1) | 0.054 (0.48) |
| **dA** | -0.0027 (0.00061) | -4.41 (131.4) | 2.1 x $10^{-5}$ **(0.0004)** |
| **A0:dA** | -0.00022 (4.3 x $10^{-5}$) | -5.13 (131.6) | 1 x $10^{-6}$ **(< $10^{-4}$)** |
| **Low Alpha Peak Frequency** | | | |
| **A0** | -0.0064 (0.002) | -3.19 (131.4) | 0.0018 **(0.021)** |
| **dA** | -0.004 (0.0032) | -1.23 (133.1) | 0.22 (0.96) |
| **A0:dA** | -0.00043 (0.00023) | -1.88 (134.1) | 0.063 (0.59) |
| **Low Alpha Band Power** | | | |
| **A0** | 0.0015 (0.0019) | 0.79 (131.1) | 0.43 (1) |
| **dA** | -0.0024 (0.0015) | -1.56 (131.5) | 0.12 (0.81) |
| **A0:dA** | -0.00025 (0.00011) | -2.29 (131.8) | 0.024 (0.3) |
| **High Alpha Peak Frequency** | | | |
| **A0** | 0.0049 (0.0018) | 2.79 (131.4) | 0.0061 (0.07) |
| **dA** | 0.0015 (0.0027) | 0.57 (132.9) | 0.57 (1) |
| **A0:dA** | 9.9 x $10^{-5}$ (0.00019) | 0.52 (133.8) | 0.6 (1) |
| **High Alpha Band Power** | | | |
| **A0** | -0.0043 (0.0014) | -3.09 (131.2) | 0.0025 **(0.027)** |
| **dA** | -0.0066 (0.0014) | -4.81 (131.8) | 4.1 x $10^{-6}$ **(< $10^{-4}$)** |
| **A0:dA** | 4 x $10^{-5}$ (9.7 x $10^{-5}$) | 0.42 (132.2) | 0.68 (1) |
| **Low Beta Peak Frequency** | | | |
| **A0** | -0.0017 (0.0063) | -0.27 (131.2) | 0.78 (1) |
| **dA** | -0.022 (0.0077) | -2.9 (132.2) | 0.0044 (0.071) |
| **A0:dA** | -0.0012 (0.00054) | -2.17 (132.7) | 0.032 (0.37) |
| **Low Beta Band Power** | | | |
| **A0** | 0.002 (0.00087) | 2.26 (131.1) | 0.026 (0.27) |
| **dA** | -0.0017 (0.00063) | -2.67 (131.4) | 0.0085 (0.12) |
| **A0:dA** | -0.00014 (4.5 x $10^{-5}$) | -3.18 (131.6) | 0.0018 **(0.028)** |
| **High Beta Peak Frequency** | | | |
| **A0** | -0.0092 (0.0059) | -1.55 (131.2) | 0.12 (0.79) |
| **dA** | -0.012 (0.0057) | -2.12 (131.8) | 0.036 (0.41) |
| **A0:dA** | -0.00022 (0.0004) | -0.55 (132.1) | 0.58 (1) |

| | | | |
|---|---|---|---|
| **High Beta Band Power** | | | |
| **A0** | 0.0005 (0.00067) | 0.76 (131.1) | 0.45 (1) |
| **dA** | -0.0024 (0.00053) | -4.55 (131.5) | $1.2 \times 10^{-5}$ **(0.0001)** |
| **A0:dA** | -0.00024 ($3.7 \times 10^{-5}$) | -6.45 (131.8) | 1.9 x 10-9 **($< 10^{-4}$)** |
| **Gamma Peak Frequency** | | | |
| **A0** | 0.0017 (0.0078) | 0.22 (133.8) | 0.83 (1) |
| **dA** | 0.045 (0.02) | 2.29 (136.6) | 0.024 (0.3) |
| **A0:dA** | 0.0011 (0.0014) | 0.75 (141.5) | 0.45 (1) |
| **Gamma Band Power** | | | |
| **A0** | $1.7 \times 10^{-5}$ (0.00015) | 0.11 (125.7) | 0.91 (1) |
| **dA** | $-6.8 \times 10^{-5}$ (0.00035) | -0.2 (127.0) | 0.85 (1) |
| **A0:dA** | $-1.3 \times 10^{-6}$ ($2.5 \times 10^{-5}$) | -0.05 (131.4) | 0.96 (1) |
